# Detecting interspecific positive selection using convolutional neural networks

**DOI:** 10.1101/2024.11.18.624117

**Authors:** Charlotte West, Conor R. Walker, Shayesteh Arasti, Viacheslav Vasilev, Xingze Xu, Nicola De Maio, Nick Goldman

## Abstract

Traditional statistical methods using maximum likelihood and Bayesian inference can detect positive selection from an interspecific phylogeny and a codon sequence alignment based on model assumptions, but they are prone to false positives due to alignment errors and can lack power. These problems are particularly pronounced when faced with high levels of indels and divergence. Leveraging the feature-detection capabilities of convolutional neural networks (CNNs), we achieve higher accuracy in detecting selection across a specific range of phylogenetic scenarios and evolutionary modes. This advantage is particularly evident with noisy data prone to misalignments. Our method shows some ability to account for these errors, where most statistical frameworks fail to do so in a tractable manner. We explore generalisability and identify future avenues to achieve broader utility. Once trained, our CNN model is faster at test time, making it a scalable alternative to traditional statistical methods for large-scale, multi-gene analyses. In addition to binary classification (inference of the presence or absence of positive selection during the evolution of the sequences), we use saliency maps to understand what the model learns and observe how this could be leveraged for sitewise inference of positive selection.

## 1 Introduction

### 1.1 Background

Natural selection is a crucial aspect of evolution. We study it to uncover evolutionary mechanisms that drive biodiversity, adaptation, and the development of life on Earth. A deeper understanding of selection allows us to make insights across biological fields, including drivers of cancer evolution (Martincorena et al., 2017), host-pathogen co-evolution (Bishop, Dean, and Mitchell-Olds, 2000) and adaptation to extreme environments (Shin et al., 2014). These insights offer practical applications, from innovative cancer treatments to effective climate change management. However, the urgency and impact of such practicalities and the growing throughput of sequencing technologies necessitates fast and accurate methods for studying selection.

Natural selection is typically studied either at the population level (microevolutionary) or at the interspecies level (macroevolutionary). Within these temporal frames, we may be interested in particular genes or genomic loci targeted by selection. In this study we focus on selection inference of protein-coding genes at the interspecies scale. This involves firstly gathering homologous genes from different species and aligning these sequences into a multiple sequence alignment (MSA). Then a phylogenetic tree that describes the evolutionary history of these genes is inferred from the MSA. Current state-of-the-art methods then take the MSA, tree and an explicit model of molecular evolution and apply statistical methods such as maximum likelihood or Bayesian inference to estimate parameters of the model that are interpreted to signal the presence or absence of selection.

Nucleotide models of sequence evolution neglect the coding and non-independent aspects of the evolution of adjacent sequence positions and lack a direct parameter through which we can model Darwinian selection of amino acid sequences. Amino acid models do not consider synonymous mutations and are largely unable to help in the detection of selection (although Weber et al., 2020, demonstrate that some information about non-synonymous mutations remains when amino acid sequence data are embedded in a genetic code-aware model). Codon-based models were proposed to address these issues. The codon substitution models proposed by Goldman and Yang (1994) and Muse and Gaut (1994) and further developed by (e.g.) Yang, Nielsen, et al. (2000) are continuous-time Markov chains defined in terms of the substitution rates from each codon to any other that can be realised by a single nucleotide change (excluding stop codons). The framework simultaneously utilises both nucleotide and amino acid-level information available in an alignment of protein-coding DNA sequences, allowing for parameters through which selection can be modelled. In particular, the parameter *ω*, frequently referred to as d N/d S and representing the ratio of the rates of non-synonymous to synonymous substitutions, has been widely used as a proxy for selective pressure (Yang, 2014). Values of ω larger than 1 suggest that substitutions leading to an amino acid change, and therefore presumably an alteration in protein structure or function, are occurring at a higher rate than synonymous (and supposedly neutral) substitutions. Typically ω > 1 is interpreted as positive selection, ω= 1 as neutral evolution, and ω < 1 as purifying selection.

Tools such as CODEML from the PAML package (Yang, 2007; Álvarez-Carretero, Kapli, and Yang, 2023) and HyPhy (Pond, Frost, and Muse, 2005) use maximum likelihood methods to infer whether natural selection affected the evolution of given protein-coding sequences through hypothesis tests and the estimation of values of *ω*. Indeed, in CODEML *ω* can be estimated for the whole tree and alignment, or per branch, per site, or both (Goldman and Yang, 1994; Muse and Gaut, 1994; Yang, 1998; Yang, Nielsen, et al., 2000; Yang and Nielsen, 2002; Zhang, Nielsen, and Yang, 2005). However, these statistical methods are prone to some issues. Firstly, they can be overly conservative (Wong, Yang, et al., 2004; Yang, Goldman, and Nielsen, 2009), resulting in the failure to detect positive selection. This is especially prevalent with sequences of very low or high divergence where the signal for positive selection is weak (Jordan and Goldman, 2012). Secondly, these methods can return false positives due to alignment errors in the MSAs (Fletcher and Yang, 2010). This is because the statistical models interpreting the MSAs do not account for misalignments and take the MSA as truth (Wong, Suchard, and Huelsenbeck, 2008). Misalignments, particularly over-alignment where non-homologous residues are placed in the same column (Löytynoja and Goldman, 2008), can result in extra inferred non-synonymous mutations and thus cause inflation of the estimated ω value.

On the other hand, misalignments can also result in false negatives. For example, if synonymous but non-homologous codons are aligned at a site under positive selection, this can lead to an apparent increased synonymous substitution rate and thus decreased *ω*. Alternatively, if truly homologous codons are not correctly aligned at a positively selected site, this may result in less supporting evidence for positive selection there. Both instances reduce statistical power at that site (Jordan and Goldman, 2012).

One way to account for alignment errors is not to assume a fixed alignment, but instead to integrate over alignment uncertainty with Bayesian methods — jointly estimating the alignment, model parameters, and the presence or absence of positive selection (Redelings, 2014). Although this approach can perform well in minimising the problem of high false positive rates (Redelings, 2014), it is computationally demanding and application is typically limited to small datasets (Pečerska, Gil, and Anisimova, 2021). Misalignment thus remains a considerable problem in selection inference analyses.

### 1.2 AI as an alternative

A promising alternative to these model-based statistical methods for the inference of selection at the interspecies level is the use of machine learning. Specifically, convolutional neural networks (CNNs) have been successfully applied to a range of problems, most notably in the field of image recognition (Krizhevsky, Sutskever, and Hinton, 2017). Since genomic MSAs have intuitive image representations, and because CNNs have an innate ability for feature detection, questions of evolution lend naturally to CNNs.

CNNs have been successfully applied in population genetics, which often relies on complex models to describe evolutionary processes affecting genetic variation within and between populations, including mutation, recombination, migration, selection, and demographic history. Consequently, calculating the likelihood of a population genetic dataset can be computationally intractable (Bertl et al., 2017). CNNs offer computationally tractable solutions to these problems. For example, Flagel, Brandvain, and Schrider (2019) use a supervised CNN to infer selective sweeps and recombination rates, where the input ‘image’ is a population genetic alignment of binary values representing allele status. ImaGene (Torada et al., 2019) uses CNNs to quantify natural selection from aligned population genomic data, with distinct alleles represented by colours.

In the context of interspecies evolutionary genetic inference, a number of machine learning approaches have recently been proposed for addressing the problem of phylogenetic tree inference (Suvorov, Hochuli, and Schrider, 2020; Zou et al., 2020; Solis-Lemus, Yang, and Zepeda-Nunez, 2022; Nesterenko et al., 2024) and phylogenetic model selection (Burgstaller-Muehlbacher et al., 2023). To our knowledge, such approaches have not previously been used for studying interspecific selection.

CNNs present a compelling alternative to traditional statistical methods for detecting interspecific positive selection. Genetic data often exhibit complex, non-linear relationships between features, challenging conventional statistical approaches. Combining feature extraction and non-linearity (obtained through layers of non-linear transformations) means that CNNs are well-equipped to deal with the intricate patterns that occur in an MSA when alignment error interacts with selection. The features learned by a CNN are based on the data it encounters during training. In the study of selection, the range and complexity of simulation models available for creating realistic training data (e.g. Sipos et al., 2011; Mallo, De Oliveira Martins, and Posada, 2016; Haller and Messer, 2019) surpass those of models tractable for maximum likelihood inference. Moreover, CNNs are highly adaptable, allowing for task- or model-specific optimisation. Training an AI model is resource- and time-intensive but, once trained, AI methods typically result in faster inference compared with maximum likelihood or Bayesian methods. This means that AI could also address the problem of excessive computational resource consumption of classical phylogenetic methods when used on large genomic datasets (see Kumar, 2022; Kozlov and Stamatakis, 2024, and references therein).

### 1.3 Introducing OmegaAI

Here we tackle the problem of inferring selection at the interspecies level while accounting for alignment uncertainty and errors. We introduce OmegaAI, a CNN trained to produce a binary classification indicating the presence or absence of positive selection given an MSA of homologous, protein-coding nucleotide sequences from different species. We aim to address the problems caused by alignment errors by training our CNN on alignments containing such errors. We show that our approach can achieve high levels of accuracy when inferring positive selection, especially at greater divergences, even with high levels of alignment error. Further, data evaluation time is approximately four orders of magnitude faster on average compared with state-of-the-art maximum likelihood methods. This work aims to be a proof of principle, so whilst the method’s application is limited to a specific set of problems, we nevertheless illustrate the promise of machine learning methods for studying selection, and discuss possibilities for future advances that may be even more effective.

## 2 New Approaches

### 2.1 CNN architecture

Following previous studies that have applied deep learning methods to evolutionary questions (Flagel, Brandvain, and Schrider, 2019; Torada et al., 2019; Suvorov, Hochuli, and Schrider, 2020), we choose a CNN model for our problem of detecting positive selection from input data which, before pre-processing, is an MSA of protein-coding, nucleotide sequences. CNNs are characterised by convolutions. The OmegaAI architecture (Figure 1) begins with a series of convolutional blocks. The first layer uses a filter size of m ⇥ 3, where m is the number of sequence rows (taxa) and 3 corresponds to one codon alignment column (i.e. 3 nucleotide or gap characters), with a stride of 3 to evaluate each codon column separately, with the intent to encode the concept of a codon in this initial layer. There are a total of six convolutional layers with ReLU activation, batch normalization, dropout for regularization, and average pooling — operations that are standard with regards to CNN architectures (De Andrade, 2019). A global pooling layer condenses the feature maps into a single vector, which is then processed by a fully connected layer. The final layer is a dense layer with a sigmoid activation function, producing a single binary prediction (selection, 1, vs. no selection, 0), indicative for the whole MSA. As a proof of principle study, our aim was to create a concise yet effective machine learning model, hence our choice to use a conservative number of layers and parameters. Once fully trained, this CNN is what we refer to as an OmegaAI model. Throughout this work, several OmegaAI models were trained under various simulation scenarios, each reflecting aspects of biological reality.

**Fig. 1.**
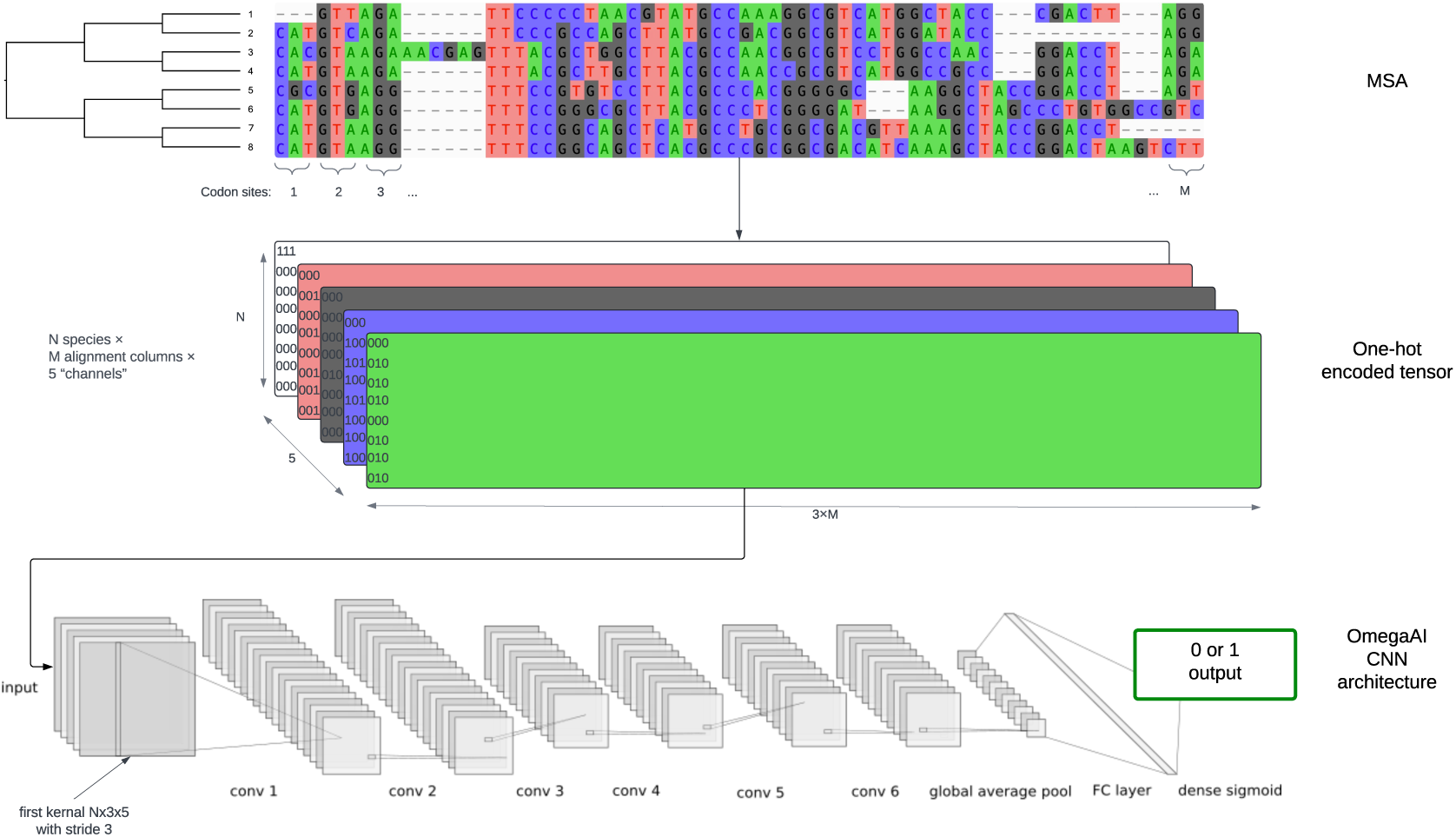
OmegaAI model architecture. After sequence evolution is simulated along an 8-taxon tree and aligned to result in an MSA (top), the MSA is converted into a one-hot encoded tensor in which each nucleotide has a binary representation in the depth dimension (middle). Batches of these tensors are fed to the CNN in order to train it. There are 6 convolution layers followed by global average pooling, a fully connected layer and finally a dense sigmoid layer, resulting in a single output of 1 (positive selection) or 0 (no selection) (bottom). Each convolution block consists of convolutions followed by batch normalisation, drop out regularisation and average pooling. CNN architecture diagram was created using NN-SVG (LeNail, 2019).

### 2.2 Simulation

In the absence of large amounts of biological data known to have been subject to positive, neutral or purifying selection, we use simulations under a variety of conditions to generate datasets — both with and without positive selection — in the quantities required to train and test our CNN. To reduce the vast parameter space of possible simulation scenarios, we restrict our attention to the evolution of a protein-coding gene sequence on a symmetric 8-taxon tree (Supplementary Fig. 1).

We simulated nucleotide sequence evolution on this phylogeny with INDELible (Fletcher and Yang, 2009), using standard codon and site variation models that allow for different purifying, neutral, or positive selective pressures (those used for inference in PAML; Yang, 2007) as well as insertions and deletions. Many different random model parameter values were used to generate alignments with varying root ancestral protein length, strength of selection, number of sites affected, and rate of evolution. By resampling parameters across numerous simulations, we generated a comprehensive training set covering a variety of realistic selective scenarios.

To further mimic real-life scenarios, we cannot rely on knowledge of the true sequence alignments but rather we must re-align the simulated sequence datasets. We primarily use Clustal Omega v1.2.4 (Sievers, Wilm, et al., 2011) for this since it is fast and widely used (Sievers and Higgins, 2018), although we also consider other alignment tools for comparison purposes. By training OmegaAI on MSAs containing alignment errors, we hope OmegaAI will learn to account for them in its predictions.

### 2.3 Training

Our aim is to train OmegaAI to produce a single binary classification for the whole MSA, stating whether or not the gene has undergone positive selection. The CNN is trained on 1,000,000 MSAs, with exposure to roughly equal amounts of genes with and without positive selection. Although in reality most genes might not have sites under positive selection, balanced training data is used to avoid bias towards the majority class (Wei and Dunbrack, 2013; Ghosh et al., 2024).

Simulated MSAs are converted into one-hot encoded tensors, a binary representation suitable for the numeric nature of neural networks, before being input into the model. During training, these tensors pass through the convolutional layers in batches, where feature extraction and transformation occur, followed by pooling layers that reduce spatial dimensions. Batch normalization and dropout layers regularise the model to prevent overfitting. The model weights are adjusted based on the loss computed from the difference between the predicted output and the true label, using backpropagation and the update rules of the Adam optimiser (Kingma and Ba, 2017). This process is repeated for multiple epochs, with each epoch exposing the CNN to the entire training dataset exactly once. A CNN requires multiple epochs to iteratively refine its weights through gradient descent, allowing it to effectively learn and generalise from the training data. The training dataset is divided into training and validation subsets to monitor performance, resulting in the final model.

### 2.4 Testing

A further 2,000 MSAs are simulated in the same manner to test each OmegaAI model, guaranteeing avoidance of train-test contamination (Magar and Schwartz, 2022). Test data analysed by the CNN is also evaluated using CODEML, and results are compared. We chose CODEML for benchmarking OmegaAI’s machine learning results as it implements a state-of-the-art maximum likelihood method for detecting selection.

Deep learning methods are often criticised for being “black boxes”. In order to gain insight into what the OmegaAI CNN learns, we visualise saliency maps (Simonyan, Vedaldi, and Zisserman, 2014) that indicate which parts of each input dataset, and in particular which alignment columns, are contributing to the prediction obtained for that dataset. We compare these findings to the sitewise inference of positive selection offered by CODEML.

An overview of the method workflow is given in Figure 2 and technical details regarding all methods are discussed in the Methods section.

**Fig. 2.**
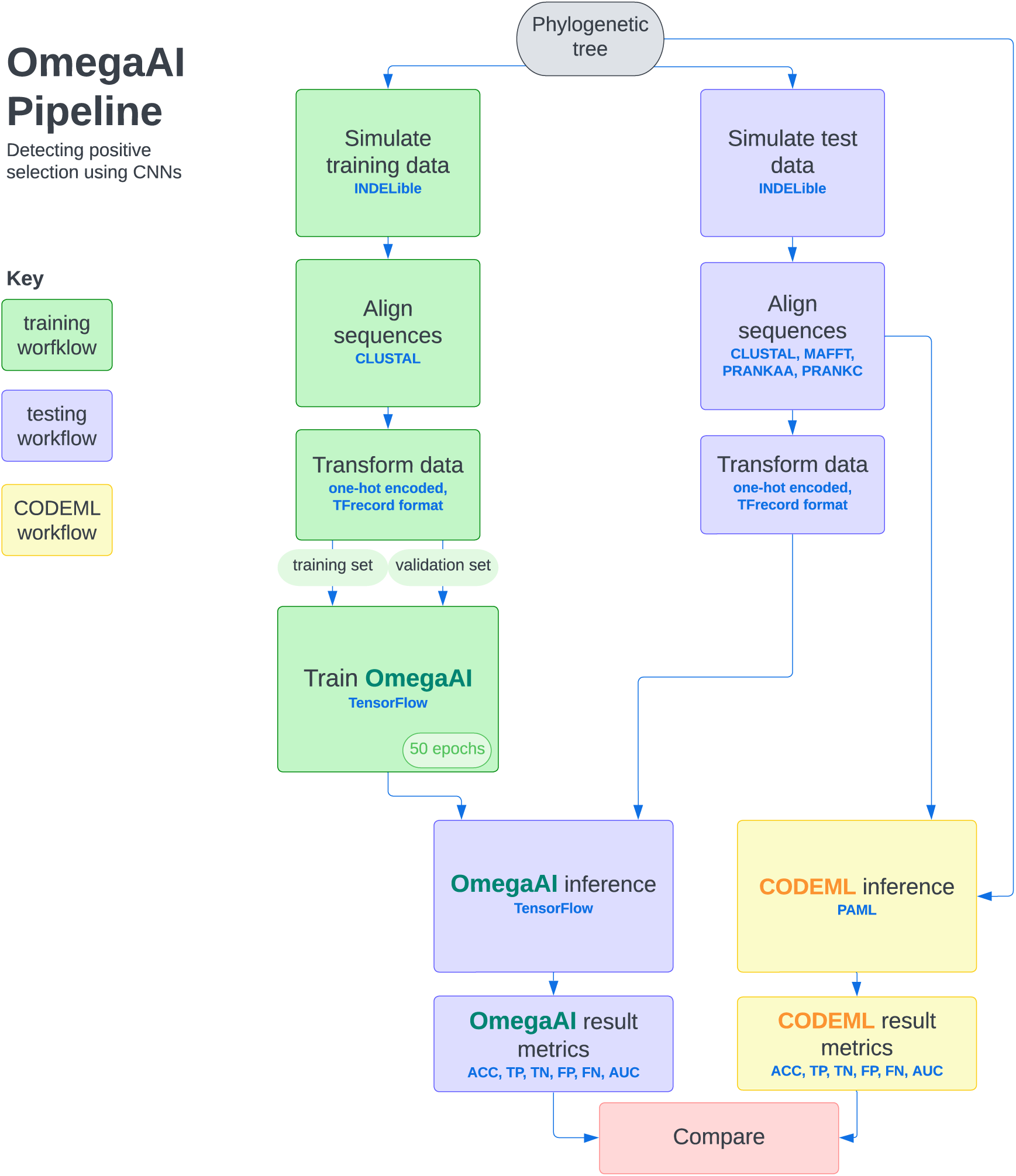
Graphical summary of the OmegaAI training and testing workflows. The workflow to train an OmegaAI model is shown in green. Sequence evolution is simulated along a phylogenetic tree under positive or purifying/neutral selection using INDELible, then typically aligned using Clustal. These alignments are one-hot encoded and collated into TFrecord format for efficient batching. The OmegaAI model is trained for 50 epochs. Sequences for the testing workflow (lilac) are simulated under the same parameters as training, and aligned using four aligners: Clustal, MAFFT, PRANKaa and PRANKc. These are also similarly transformed for testing the OmegaAI model, or passed directly in fasta format to CODEML (yellow). Binary classifier metrics ACC, TP, TN, FP, FN, AUC are calculated for both methods and then compared. See main text for full details.

## 3 Results

### 3.1 OmegaAI performs well with baseline parameters

We choose the symmetric 8-taxon tree to be the known, true phylogeny (Supplementary Fig. 1) along which sequence evolution is simulated. Our baseline parameter set describes a tree scaling of 0.2 substitutions per site per branch, with an indel rate of 0.1 per substitution. This choice serves as our reference point for analysis, chosen based on literature and empirical data (Ogurtsov, Sunyaev, and Kondrashov, 2004; Fletcher and Yang, 2010; Jordan and Goldman, 2012), and due to the fact that these parameters produce realistic alignments (Supplementary Fig. 2 and Supplementary Fig. 3) with the divergence being high enough that we obtain a reasonable signal of positive selection when it is present, without encountering the excessive information loss and alignment error that we see with higher divergences. Along this phylogeny, data is simulated with and without selection with parameters as described in detail in Methods. OmegaAI is trained using 1,000,000 MSAs, with performance of trained models assessed using 2,000 independently simulated MSAs (see New Approaches and Methods for details).

To obtain binary classifications from real-valued data, it is necessary to determine thresholds that delineate the categorical boundaries between positive selection and no selection. These decision thresholds influence the trade-off between precision and recall, allowing for optimisation of a preferred metric. In statistical methods like CODEML, a significance level is set to control the Type I error rate. For instance, a 95% confidence level is often chosen to detect positive selection, ensuring a false positive rate of 5% based on true alignments (Wong, Yang, et al., 2004) (see Methods). This approach provides a statistically grounded framework for decision-making.

In contrast, OmegaAI lacks an intrinsic statistical framework to establish a decision threshold. Therefore, we have selected 0.5 as a natural threshold, representing the midpoint of the output value range between 0 and 1. Unlike CODEML, which aims to control its error rate directly through its significance level, OmegaAI’s performance is evaluated using various metrics, and its error rates are empirically determined. The flexibility of adjusting the decision threshold in OmegaAI allows for optimisation based on specific application needs, but it also requires thorough benchmarking to identify the most appropriate threshold. Some applications may prioritise specificity over recall or vice versa.

To assess and compare the efficacy of detecting positive selection by OmegaAI and CODEML, in Figure 3 we show Receiver Operating Characteristic (ROC) curves which illustrate the trade-off between true positive and false positive rates when varying the methods’ decision thresholds between 0 and 1. Precision-recall curves are also presented, showing the trade-off between these metrics as decision thresholds are varied. For both metrics, the total area under the curve (AUC) provides a comprehensive evaluation of the method’s performance, encompassing all potential classification thresholds. Figure 3 A–B compares the performance of OmegaAI and CODEML when evaluated with test data aligned using Clustal. The OmegaAI curve sits above the CODEML curve in both, showing that the performance of OmegaAI dominates that of CODEML over the whole threshold range. The higher AUC value for OmegaAI in both the ROC and precision-recall plots provides a summary of this superior performance. Dots on the curves represent results obtained at typical thresholds. For example, comparing OmegaAI with a decision threshold of 0.5 with the 0.95 pvalue threshold commonly used in CODEML, we see that OmegaAI has a higher true positive rate (TPR) and lower false positive rate (FPR) (Figure 3 A) and higher precision and recall (Figure 3 B) than CODEML.

**Fig. 3.**
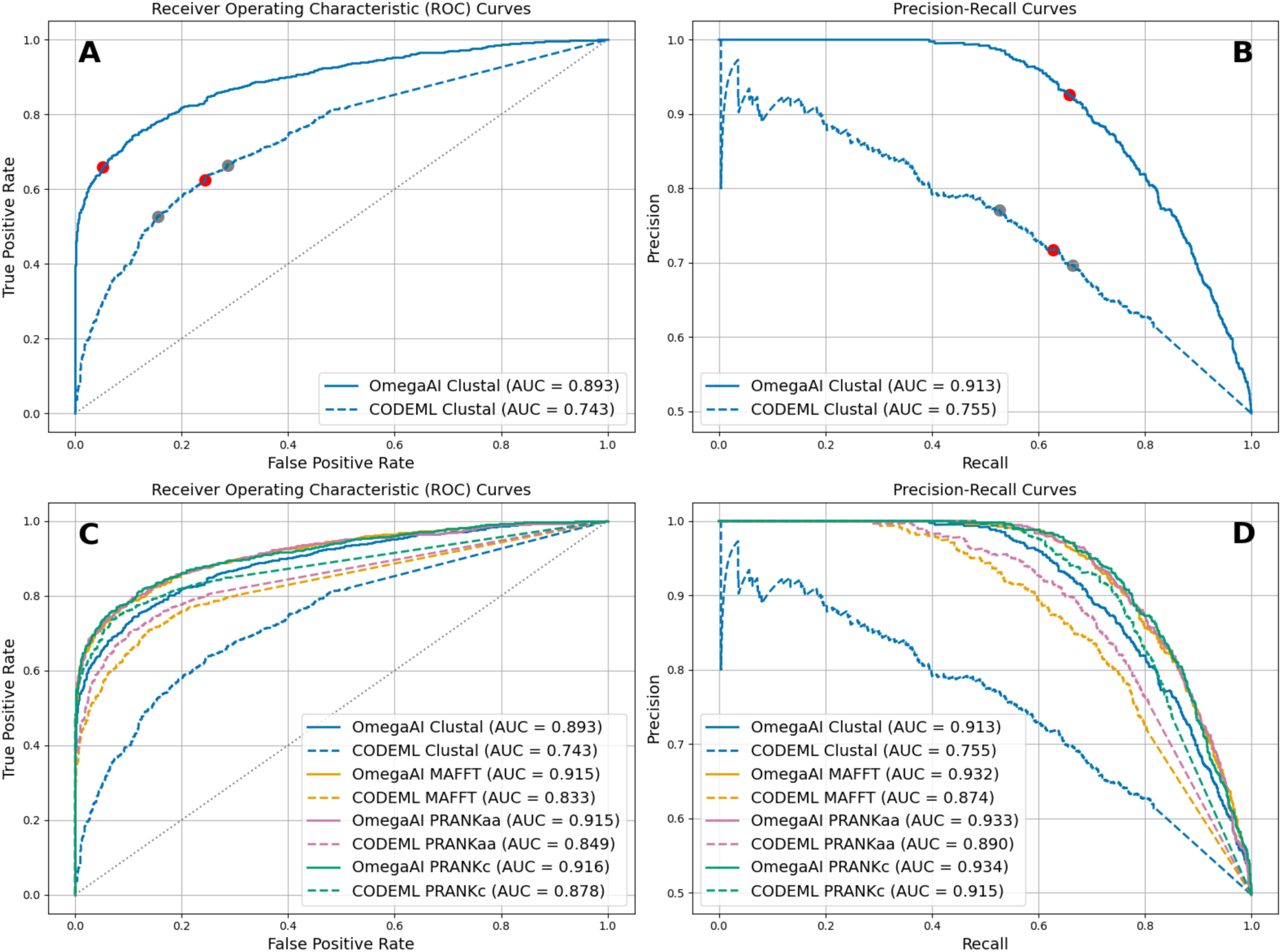
OmegaAI vs. CODEML ROC and precision-recall curves for baseline parameters. **A:** ROC curves for OmegaAI (solid line) and CODEML (dashed). The red dot on the solid line represents the OmegaAI result at the 0.5 decision threshold used throughout this work. The red dot on the dashed line represents the 0.95 p-value threshold typically used for determining positive selection with CODEML. The grey dots indicate results for p-value thresholds of 0.99 (left) and 0.9 (right). **B:** Precision-recall curves for OmegaAI (solid) and CODEML (dashed). Red and grey dots are as in (A). **C:** ROC curves when the same OmegaAI model as in (A–B), trained exclusively on Clustal alignments, is used to evaluate simulated test data that has been aligned with four different aligners: Clustal (as in A–B), MAFFT, PRANKaa and PRANKc. This test data is evaluated by both OmegaAI (solid lines) and CODEML (dashed lines). **D:** As in (C), now showing precision-recall curves.

In Figure 3 C–D we repeat this analysis using test datasets aligned with three alternative aligners, again evaluating each set with both OmegaAI and CODEML. This approach allows us to assess the performance of the selection detection methods across various alignment algorithms and qualities, and to evaluate OmegaAI’s ability to generalise to MSAs aligned by algorithms it did not encounter during training. OmegaAI consistently outperforms CODEML based on both the ROC and precision-recall metrics, for every aligner (solid lines for OmegaAI always above corresponding dashed lines for CODEML, with each colour pair representing analysis with a different aligner), as summarised by the AUC values.

The results show that OmegaAI performs similarly on PRANKc, PRANKaa and MAFFT alignments, whilst exhibiting lowest performance for Clustal alignments. Whilst the model was trained exclusively on Clustal alignments, the alignments from the other three aligners tend to be of significantly higher quality (as shown in Supplementary Fig. 2 and Supplementary Fig. 3). This suggests that alignment quality is more important for accurate prediction by our method than the CNN model learning alignment error patterns specific to any one particular alignment algorithm. Further, it appears OmegaAI’s learned ability to account for alignment error applies across alignment types.

For CODEML, there is a clearer ordering of performance by alignment type: PRANKc giving best results, followed by PRANKaa, MAFFT and then Clustal proving most difficult. This outcome agrees with the observed alignment quality rankings in Supplementary Fig. 2 and Supplementary Fig. 3. Since CODEML cannot compare data across alignment columns, more alignment errors will lead to greater information loss regarding the occurrence of selection. In particular, PRANKc and PRANKaa, as evolutionary-aware aligners, produce higher quality alignments (Jordan and Goldman, 2012) resulting in more reliable CODEML inferences.

### 3.2 OmegaAI performs well with increasing divergence and higher indel rates

We repeated this analysis across different scalings of the simulation tree, representing different divergence levels among the taxa. This approach tests simulated sequences of genes evolving over a range of rates or times. Greater divergence reduces sequence similarity and can make alignment and accurate selection signal extraction more challenging due to the increased noise.

We present 10 OmegaAI models, each trained on one of 10 datasets. The topology of the tree underlying sequence evolution simulation is unchanged, but each training dataset has incrementally increasing divergence for the branches (0.1, 0.2, …, 1.0 substitutions per codon site per branch). Likewise, we employ 10 test datasets, each exhibiting the same incremental levels of divergence to test each corresponding OmegaAI model.

We compare OmegaAI and CODEML across increasing divergences in Figure 4. We show results for test data aligned with the four different aligners, as previously. Both OmegaAI and CODEML show superior predictive power for detecting positive selection at lower divergences compared to higher ones. However, OmegaAI achieves higher accuracy and TPR compared to CODEML, whilst maintaining low FPR, across most of the aligner-divergence combinations tested.

**Fig. 4.**
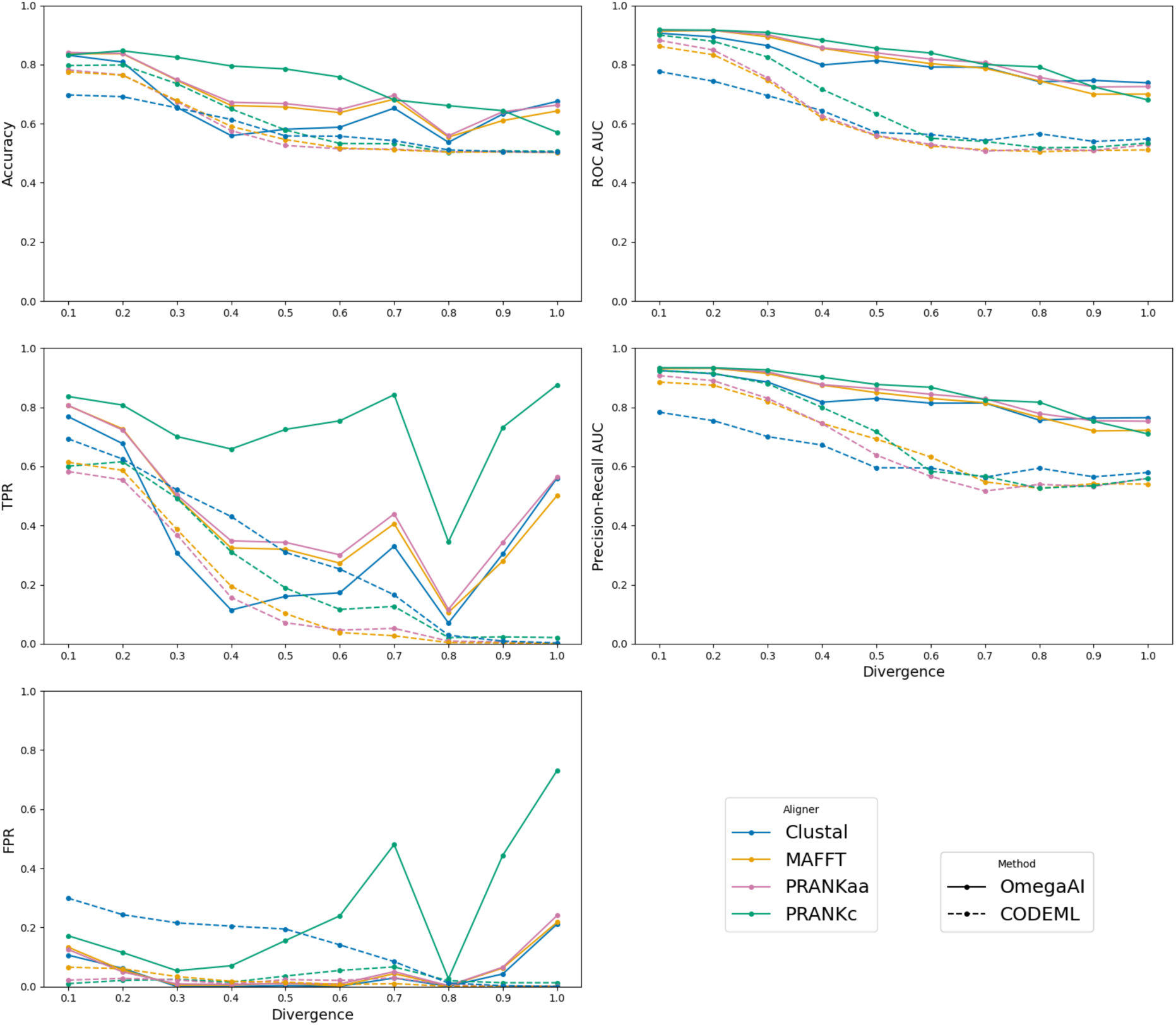
OmegaAI vs. CODEML across divergences. Various binary classifier performance metrics are presented to compare the two methods. In the left column the accuracy (defined as the number of correct predictions divided by the total number of predictions), TPR and FPR are calculated using thresholds of 0.5 and p=0.95 for OmegaAI and CODEML, respectively. In the right column AUC values for ROC and precision-recall are presented. The x-axis refers to the divergence scaling of branches of the 8-taxon symmetric tree (Supplementary Fig. 1) used for simulation. The baseline indel rate, 0.1, is used throughout. Each OmegaAI model is individually trained for each divergence using Clustal alignments. The test data is aligned by the four different aligners indicated and tested by both methods. Baseline results that were shown in Figure 3 are therefore now points shown at the relative divergence of 0.2.

CODEML’s predictive power falls with increasing divergence, as evidenced by TPR and FPR that converge towards zero. OmegaAI detects selection with TPR above zero and generally produces higher accuracies, but makes errors as evidenced by having slightly higher FPR at higher divergences, with PRANKc results being an exception of considerably higher FPR. Since OmegaAI’s AUC values are well above 0.5, we could choose to adjust decision thresholds to reduce FPRs whilst sacrificing some power, in situations where precision is prioritised over recall.

Evaluating ROC and precision-recall AUC data shows a method’s performance independent of the choice of any particular threshold. OmegaAI produces AUCs greater than CODEML across the range of divergence and aligners that we test. These results indicate that OmegaAI’s performance can be very good, comparable and frequently better than state-of-the-art maximum likelihood methods in CODEML, in various use-cases, whether emphasizing maximal true positive recovery, high precision, or minimizing false positive rates, contingent upon selecting an appropriate threshold.

Where OmegaAI has been trained using Clustal alignments, we hypothesise that the model is able to account for some amount of misalignment, and perhaps even draw information from this (see Discussion). Despite not being trained on the other aligners, OmegaAI remains able to achieve higher accuracy when evaluating these alignments than evaluation of Clustal alignments. In contrast, CODEML performs particularly poorly on Clustal alignments. This is likely due to the relatively high amount of misalignment that one can expect from Clustal alignments (Supplementary Fig. 2 and Supplementary Fig. 3) driving up its FPR.

We note the unusual behaviour at divergence 0.8, where there is a dip in OmegAI’s TPR across aligners, interrupting the otherwise smooth trend. We suspect that this is an artefact arising from an unidentified feature of the Clustal alignment algorithm, since only Clustal alignments are used to train the CNN but the artefact is observed when analysing data aligned by all aligners studied. This theory is corroborated by Supplementary Fig. 4 and Supplementary Fig. 5, since we do not witness this behaviour on these models where OmegaAI has been trained on other alignment types.

Reduced alignment quality makes the challenge of detecting positive selection harder (Jordan and Goldman, 2012), especially for models that do not account for alignment error. Over the range that we consider, increasing divergence between sequences tends to increase the difficulty of the alignment problem and thus the selection inference problem. Another challenge to aligners that we consider is higher rates of indels. We repeated the same analysis as in Figure 4 but increased the indel rates from 0.1 to 0.2 and 0.3. Overall performance decreases for both OmegaAI and CODEML, as expected, with more variable trends (Supplementary Fig. 6 and Supplementary Fig. 7). However, we again see that OmegaAI generally outperforms CODEML across the binary classifier metrics for various alignment algorithms and tree divergence scalings at these higher indel rates.

### 3.3 Training using PRANKc alignments

OmegaAI models presented so far have been trained on alignments inferred by the fast, widely used Clustal aligner. However, as noted above, Clustal can be relatively error-prone compared to other aligners whereas PRANKc is a highly accurate alignment tool which integrates phylogenetic construction and information into its alignment algorithm (Löytynoja and Goldman, 2008) but is orders of magnitude slower than Clustal. By training OmegaAI models with PRANKc alignments and comparing their performance with OmegaAI models trained with Clustal alignments, we can explore the effects of training with alignments produced from different algorithms and assess how this affects performance, including generalisability to other alignment types. With PRANKc’s higher accuracy coming at the cost of orders of magnitude longer runtimes than Clustal, its use for model training becomes very costly. Nevertheless, extensive resources were expended to test its value.

For clarity, in this section we refer to the models trained on Clustal or PRANKc alignments as OmegaAI(Clustal) and OmegaAI(PRANKc), respectively. Training sequences for both were simulated under baseline conditions (divergence 0.2, indel rate 0.1). Both models were tested with the standard test dataset used previously, where 2,000 MSAs are realigned by each of our considered aligners: Clustal, MAFFT, PRANKaa and PRANKc. ROC and precision-recall curves from this analysis are shown in Figure 5 (with cross-divergence/aligner results in Supplementary Fig. 4, and for comparison to CODEML see Figure 3 and Figure 4). The curves, and in particular the AUC values, show that both OmegaAI(Clustal) and OmegaAI(PRANKc) perform well, and to a similar level, when tested on MAFFT, PRANKaa and PRANKc alignments. However, OmegaAI(Clustal) achieves notably higher AUC values for Clustal alignments than OmegaAI(PRANKc).

**Fig. 5.**
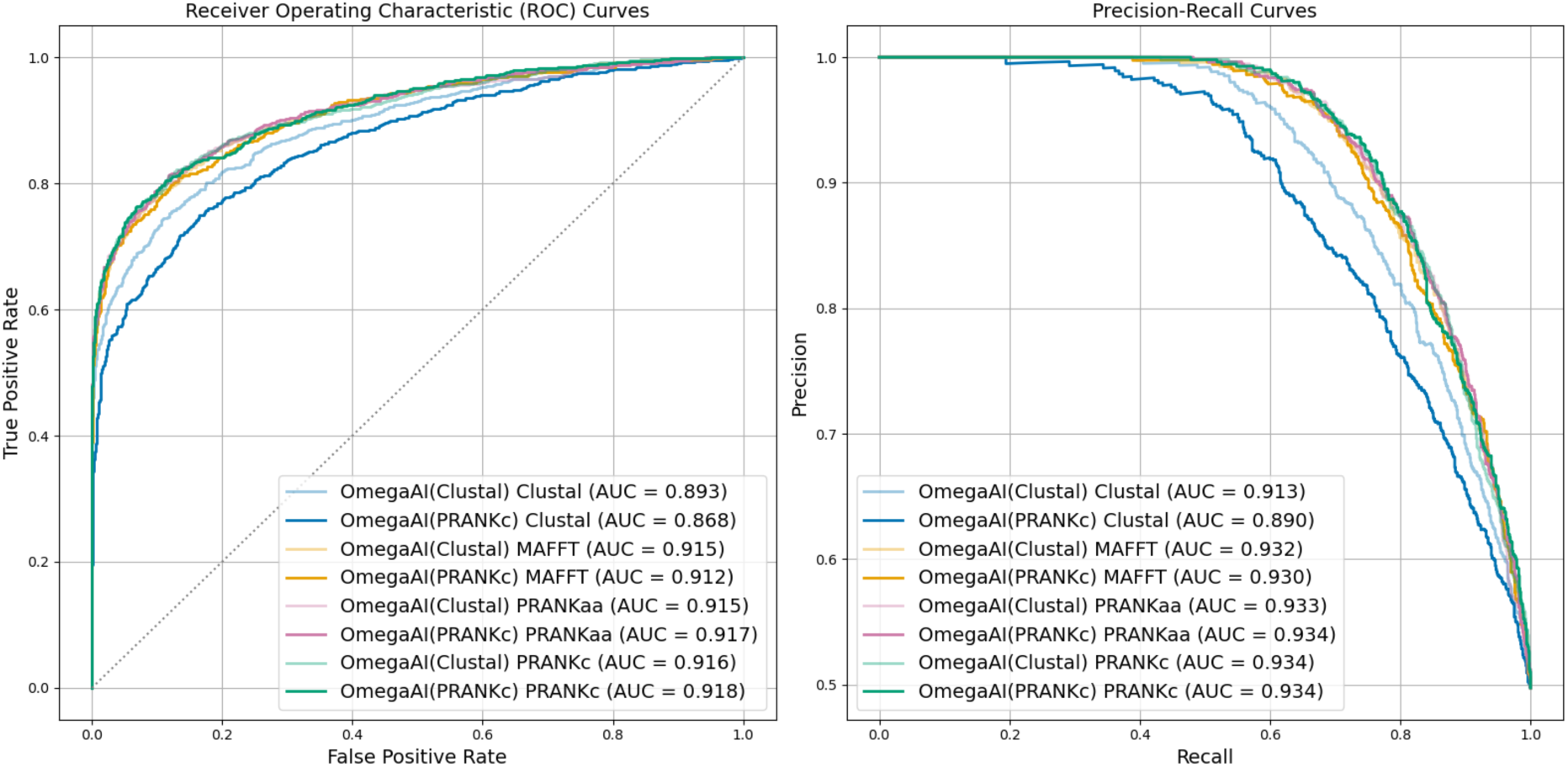
OmegaAI(PRANKc) vs. OmegaAI(Clustal). Training data was simulated under baseline parameters for both models. See main text for details of training regimes for OmegaAI(PRANKc) vs. OmegaAI(Clustal). Both models were tested on the same test data, which were aligned using four aligners. ROC (left) and precision-recall (right) curves are shown to illustrate the methods’ performance.

These findings indicate that precise alignment is not imperative during training to enhance our models’ predictive capacities, and that training on error-prone alignments makes the model more robust to alignment errors. However, during testing, alignment accuracy does significantly influence predictive accuracy, with superior results typically associated with higher-quality alignments, irrespective of training data alignment accuracy. This is evidenced by baseline tests performed on higher quality alignments producing higher AUC values than for Clustal alignments, which are less accurate (see above). Supplementary Fig. 4 illustrates a similar trend across all divergence levels studied, with OmegaAI(Clustal) generally performing better on Clustal alignments than OmegaAI(PRANKc). While OmegaAI(Clustal) excels with Clustal alignments, OmegaAI(PRANKc) occasionally outperforms OmegaAI(Clustal) on PRANKc alignments for certain divergences, with comparable performance observed for MAFFT and PRANKaa alignments when tested with the two models. These results suggest that OmegaAI models may benefit from more exposure to alignment error during training.

### 3.4 Benchmarking with the true alignment

CODEML applied to true, simulated alignments represents a gold-standard for positive selection inference because the true alignment contains no errors, artefacts or other features that CODEML’s models do not account for. The true alignment and the simulation tree should permit a highly accurate inference of the evolutionary events leading to a given set of protein coding sequences. Results from the following analysis are likely unattainable in any real-life case, since data will never conform exactly to these generative models or be perfectly aligned. However, they can help us better understand what factors contribute to the performance of different methods.

For comparison with CODEML’s inference of positive selection on true alignments we trained a new OmegaAI model solely on true simulated alignments across our divergence set. We also took the models previously trained on Clustal alignments and tested all of these on true alignments. In Figure 6 we show the ROC and precision-recall results from baseline parameter simulations (results for other divergence levels are shown in Supplementary Fig. 5). All three methods perform comparably well on the true alignment baseline test dataset, with very similar AUCs for both ROC and precision-recall. CODEML produces slightly better TPR at lower FPR than both OmegaAI methods, while the OmegaAI model trained on true alignments achieves the highest ROC AUC. These results further support the idea that the superior performance of OmegaAI in previous analyses is attributable to its ability to account for alignment errors.

**Fig. 6.**
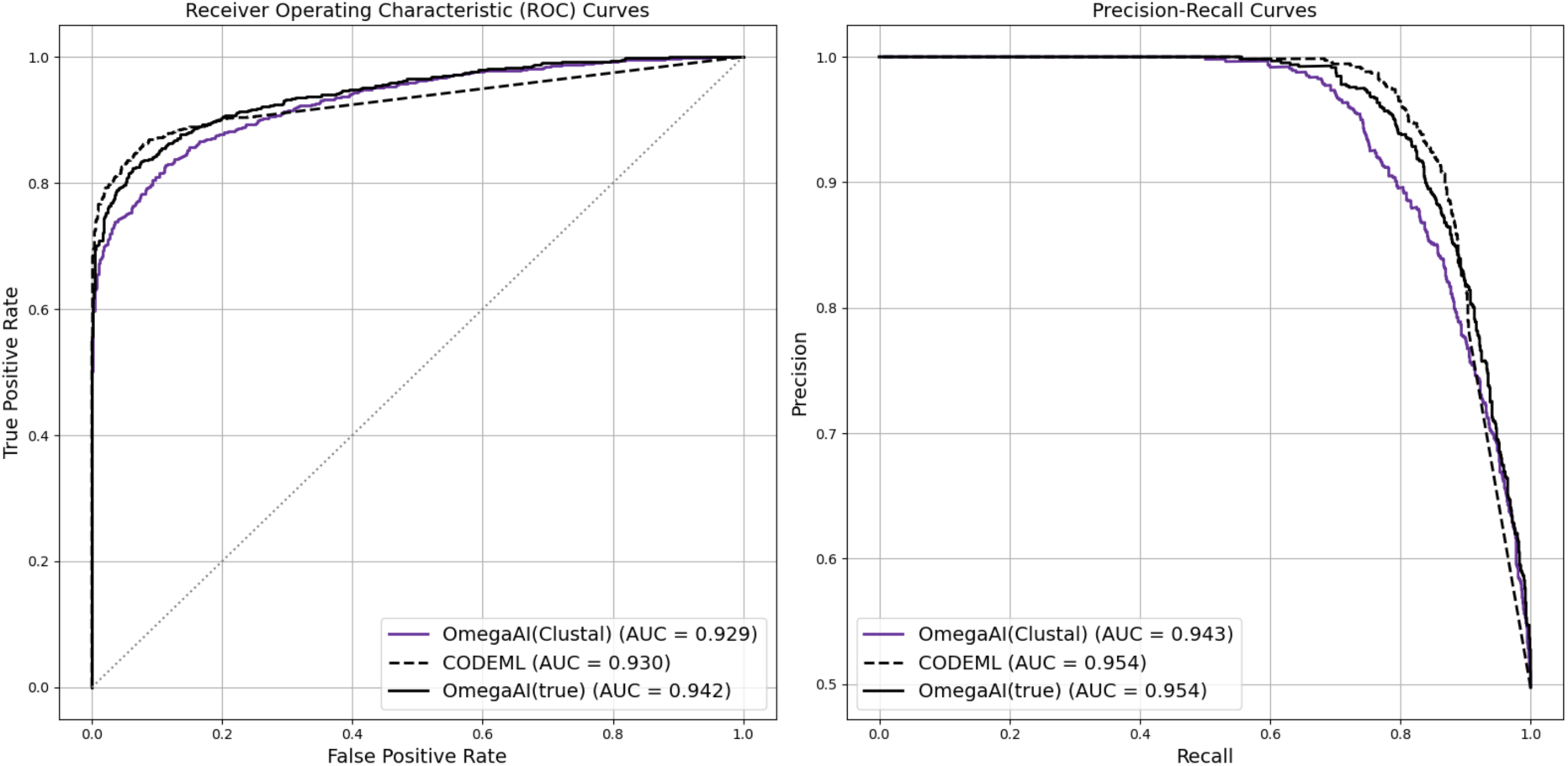
Analysing true alignments. ROC and precision-recall curves are shown to illustrate different methods’ performances. Lines labelled ’OmegaAI(Clustal)’ refer to the OmegaAI model that has been exclusively trained on Clustal alignments. ’OmegaAI(true)’ indicates training was on true alignments. Both are trained using data simulated under our baseline parameters. Both CNN models and CODEML are tested using true alignments, where the same test dataset is used for all methods.

So far when evaluating the performance of OmegaAI in terms of accuracy, TPR and FPR, we have used a decision threshold of 0.5 as a natural choice to make inferences on weights between 0 and 1 being output by OmegaAI. This choice leads to desirable results in some cases, but could be adjusted for others. For example, in Figure 4 we see that when OmegaAI evaluates Clustal, MAFFT or PRANKaa alignments, the accuracy is high whilst maintaining low FPR, compared to CODEML, for most divergences. However, whilst the accuracy of results from OmegaAI applied to PRANKc alignments are some of the highest, so too are the FPRs. We observe a similar trend when OmegaAI (trained on either Clustal or true alignments) evaluates true alignments (see Supplementary Fig. 5). From these observations, it appears that higher quality MSAs may benefit from being evaluated by OmegaAI using a lower decision threshold.

### 3.5 Generalisability

We have shown that OmegaAI models trained on Clustal alignments generalise effectively to the other aligners we have tested. However, up to this point our CNNs have been trained and tested on specific trees and divergence levels. In real-world scenarios, the data will be more varied.

AI methods can be difficult to generalise without very large and diverse training data (Eche et al., 2021). Whilst there exist strategies to mitigate this, such as fine-tuning and domain adaptation (Ganin and Lempitsky, 2015), there is a common struggle between creating models which perform well at a specific task but fail to generalise outside of it, versus models that offer wide general use but whose performance falls short for any specific subgroup (Eche et al., 2021; Huisman and Hannink, 2023). We found it valuable to consider whether OmegaAI as already studied was able to cope with a greater diversity of use cases. Here, we investigate its performance for different divergences of the 8-taxon simulation tree and in the absence of information of the underlying tree topology.

#### Training OmegaAI with reduced phylogenetic information

So far, all simulated MSAs have had the same row ordering relative to their underlying tree. As a result, the CNN is injected with learnable information about the relative relatedness of the sequences in the MSA during training (in addition to what it can learn from sequence similarity, indel patterns, and the consistent use of the same simulation tree). This structure is also predictable at test time, since the same tree and row ordering is used for the test data. Using a consistent tree and alignment row ordering during training and testing is akin to how we give CODEML a pre-estimated tree topology while testing for selection. We sought to test how well OmegaAI would perform with reduced information regarding the underlying simulation tree. This was achieved by randomly shuffling the row ordering of all MSAs before use.

Results for OmegaAI trained and tested on baseline parameters, with row shuffling, referred to here as OmegaAI(shuffled), are shown in Figure 7 (with cross-divergence/aligner results in Supplementary Fig. 8). Removing any information about an underlying tree during training negatively impacted OmegaAI(shuffled)’s prediction power, reflected by the lower ROC and precision-recall curves and reduced AUC values for the OmegaAI model trained on shuffled data. However, OmegaAI(shuffled) still outperforms CODEML by these same metrics. For some mid-range divergences and aligners the accuracy achieved by OmegaAI(shuffled) is similar to that of OmegaAI when given consistent row ordering at training and test time (Supplementary Fig. 8). However, this OmegaAI(shuffled) result, whilst exhibiting a higher TPR, comes with a dramatically increased FPR for all divergences, especially for tests performed on PRANKc alignments (a case where adjusting the decision threshold to lower the FPR and sacrifice some power could be valuable). Overall, these results suggest that the tree information is important to OmegaAI’s ability to predict and that indeed the model learns some representation of the underlying tree topology when MSA row ordering contains information. Further, it is evident that OmegaAI models, as currently trained, would likely not generalise well to a range of tree topologies to a point where they would compete with the models trained for a specific topology.

**Fig. 7.**
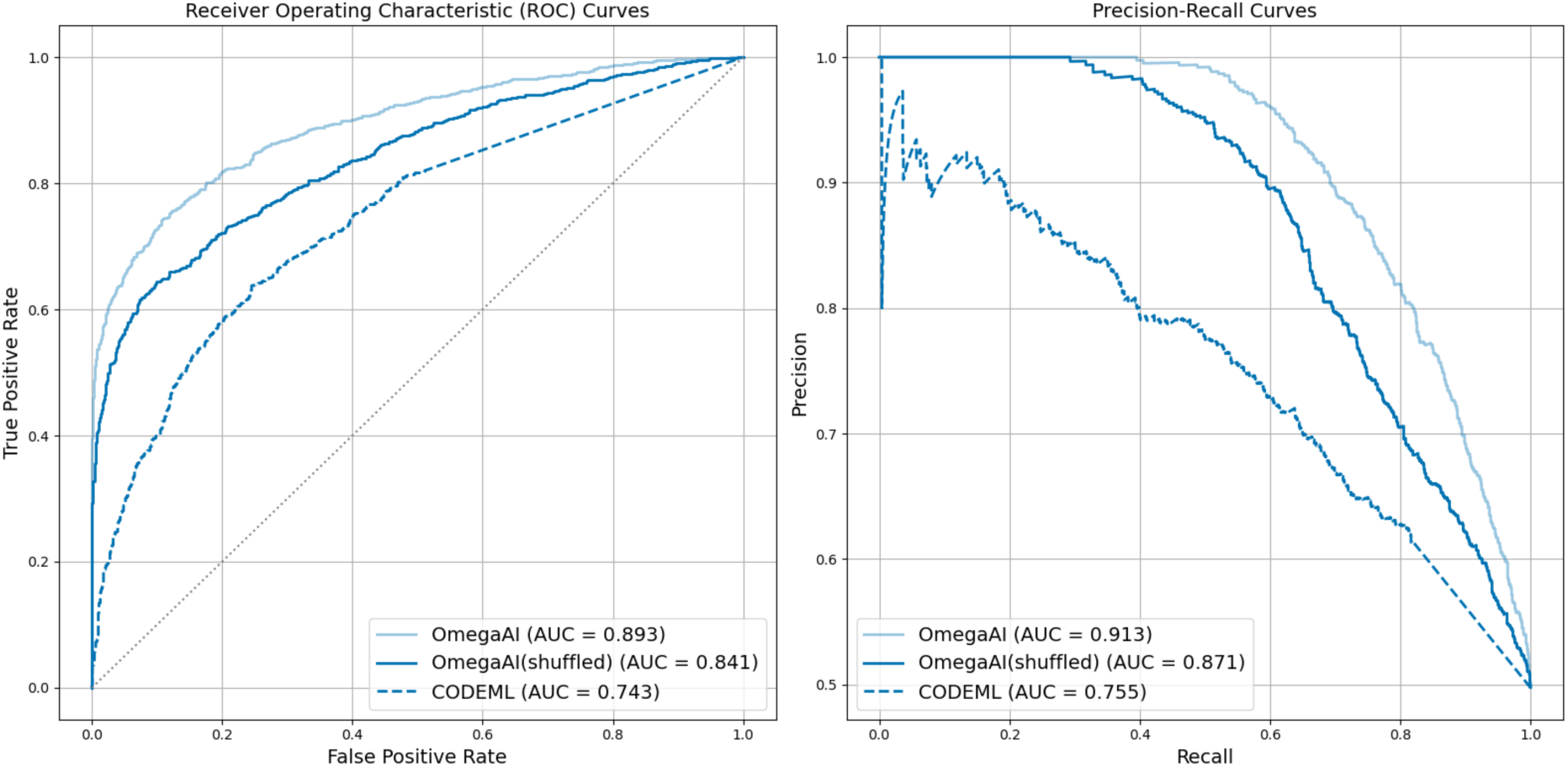
Comparing OmegaAI models with and without row shuffling. CNN models are trained with baseline parameters (divergence 0.2, indel rate 0.1). In OmegaAI(shuffled), MSA rows are randomly shuffled in both training and test data to remove some tree topology information. ROC (left) and precision-recall (right) curves are shown to illustrate the methods’ performance. CODEML represented by dashed lines for comparison; OmegaAI and CODEML results are as in Figure 3.

#### Training and testing across divergence levels

We also tested generalisability with respect to the divergence parameter of the simulation tree. By keeping the simulation tree and row ordering constant, the model can learn a representation of the topology of the sequences’ evolutionary relationships, but the rate of evolution will vary: challenging the model to deal with different levels of signal from mutations and noise from misalignment. We describe two experiments that tested this.

We tested the OmegaAI model trained on the central value (0.5) of our divergence range on data from all divergences in our set, with other parameters held constant. The results are shown in Figure 8 (left). In terms of accuracy, OmegaAI produces poor scores when tested on trees other than those with branch lengths of 0.5 (the value for which it was trained). There is a tendency in these cases for OmegaAI to infer all datasets with a lower divergence than it was trained with to have undergone positive selection, and all datasets with higher divergence to have undergone neutral/purifying selection. Comparison with an analogous approach in CODEML is shown in Supplementary Fig. 9, where CODEML also performs poorly when the assumed divergence level differs from what is being tested. However, ROC AUC values from this analysis indicate that the OmegaAI model trained on 0.5 divergence trees still effectively ranks instances of positive and negative selection for some divergences around 0.5, but the default threshold is not optimal for classifying these other divergence sets.

**Fig. 8.**
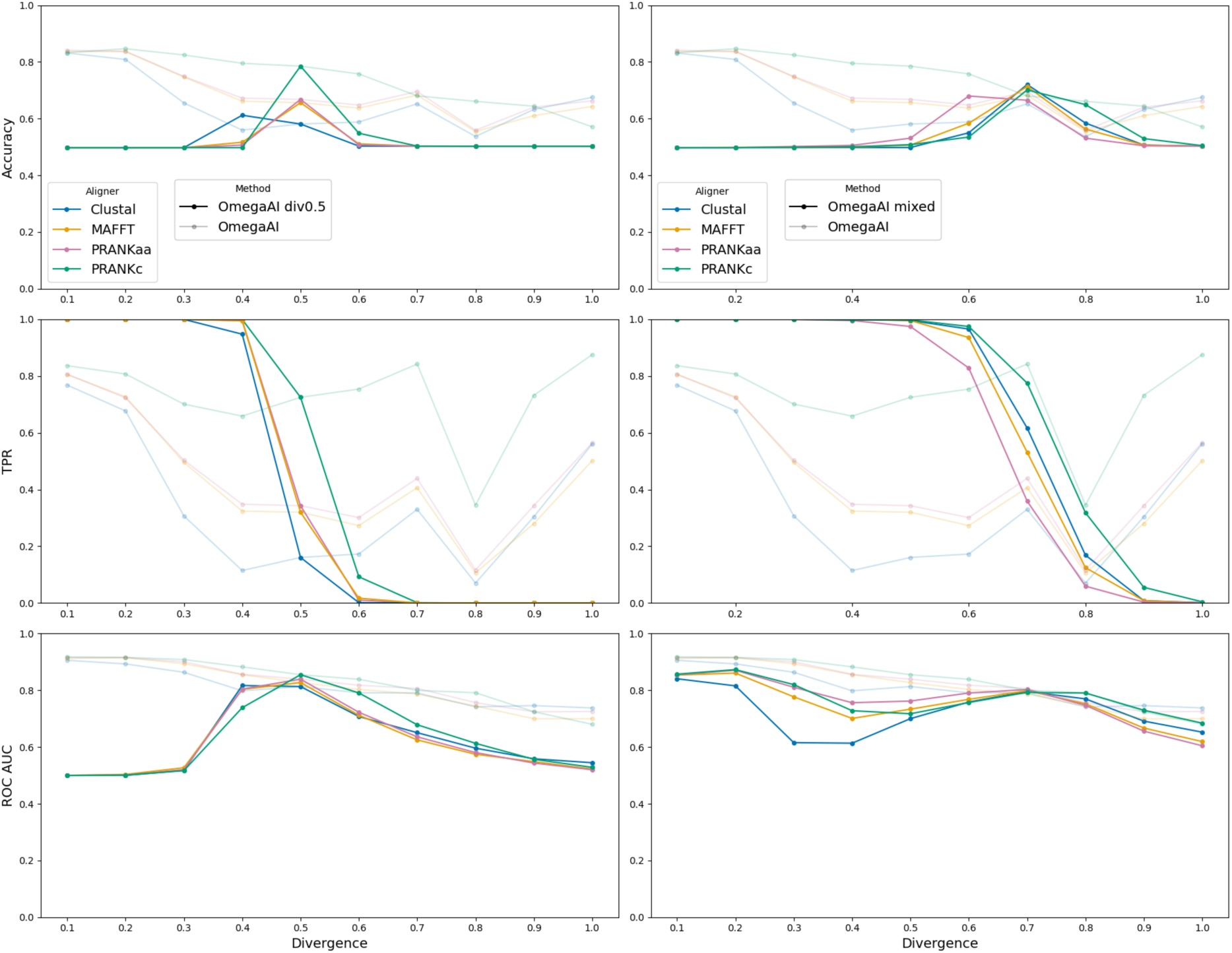
OmegaAI divergence generalisation. Left column: The single OmegaAI model trained with trees of divergence scaling 0.5 is tested on data from trees of each divergence level in the set (bold lines), and compared with multiple standard OmegaAI models retrained and tested for each individual divergence, as in Figure 4 (semitransparent lines). The accuracy and TPR are computed using a decision threshold of 0.5. ROC AUC gives a measure of performance across possible thresholds. **Right column:** A single OmegaAI model trained on 1,000,000 trees containing an equal number from each divergence level in the set. It is tested across divergences (bold) and compared to the standard OmegaAI model retrained and tested for each divergence level (semi-transparent). The accuracy and TPR result from a decision threshold of 0.5 and ROC AUC is shown to illustrate performance independently of threshold choice.

Next we created a model exposed to data from all of the 10 divergence levels in our divergence set during training, to see if this increased OmegaAI’s ability to generalise to other divergences without having to optimise for different thresholds in different testing scenarios. The training dataset was kept the same size (1,000,000 trees), but now consisting of equal numbers from each of the 10 divergences. This creates a single OmegaAI model which has learnt from phylogenies with a range of divergence levels, not just a single divergence level as with all models previously described. We tested this model on each test dataset, each simulated under a single divergence. Results are shown in Figure 8 (right).

The model exhibits similar behaviour to our previous experiment where the model trained on 0.5 divergence trees was tested across the range of divergence test sets. The difference in this case is that this new model shows reasonable performance at divergences 0.7 and 0.8, indicated by accuracy values similar to when an OmegaAI model is trained solely on trees of a single divergence level and tested on trees of the same divergence. Compared to the model trained on 0.5 divergence trees, the mixed divergence model demonstrates broader applicability across the divergence set, as indicated by its higher accuracy and especially its AUC values. This suggests that the mixed divergence OmegaAI model has learned to detect positive selection across a range of divergences, with its best performance centered at the 0.7 level, in contrast to the 0.5 model’s much narrower effective range. The TPR and AUC values together indicate that the decision threshold is too low for lower divergences and too high for higher divergences and it happens to be the case that the default threshold of 0.5 is optimal around divergences 0.7 and 0.8. More results from this analysis are shown in Supplementary Fig. 10.

In these two analyses, where accuracy is low but the ROC AUC value is good, it is evident that there is a thresholding issue. While the need to optimise thresholds in each testing scenario indicates that the OmegaAI models are not immediately generalisable, it also demonstrates that the information learned by OmegaAI in one scenario is transferable to others.

### 3.6 Alignment saliency: visualising and quantifying what OmegaAI learns

Saliency in the context of CNNs applied to image recognition tasks refers to the identification and visualisation of the most relevant pixels of an input image that contribute to the model’s prediction (Simonyan, Vedaldi, and Zisserman, 2014). It has been used as a strategy to aid understanding of what a neural network learns. In our case, where the input is an MSA instead of an image, a saliency map highlights the alignment positions that were most influential in OmegaAI’s prediction for a given MSA. Saliency is computed by taking the gradient of the output with respect to the input: the magnitude of the gradient shows which characters need the least change to most affect the output, and these can be visualized in a saliency map where brighter regions highlight key positions. For more details, see Methods.

In Figure 9 we present an example saliency map for an MSA with sequences simulated under baseline conditions with positive selection and aligned using Clustal. In such an alignment, each site evolves under purifying, neutral or positive selection, with most sites evolving under purifying or neutral selection (see Methods for details). Alignment sites are coloured according to their selection class in Figure 9, but note that more than one colour can occur in an alignment column due to misalignment. In this example, sites evolving under positive selection have *ω* = 4.28. Figure 9 also shows CODEML’s sitewise predictions of positive selection, inferred using a Bayes empirical Bayes (BEB) approach (see Methods for details). The figure reveals that high saliency tends to correlate with sites evolving under positive selection, indicating that the OmegaAI model finds these sites most informative to its predictions.

**Fig. 9.**
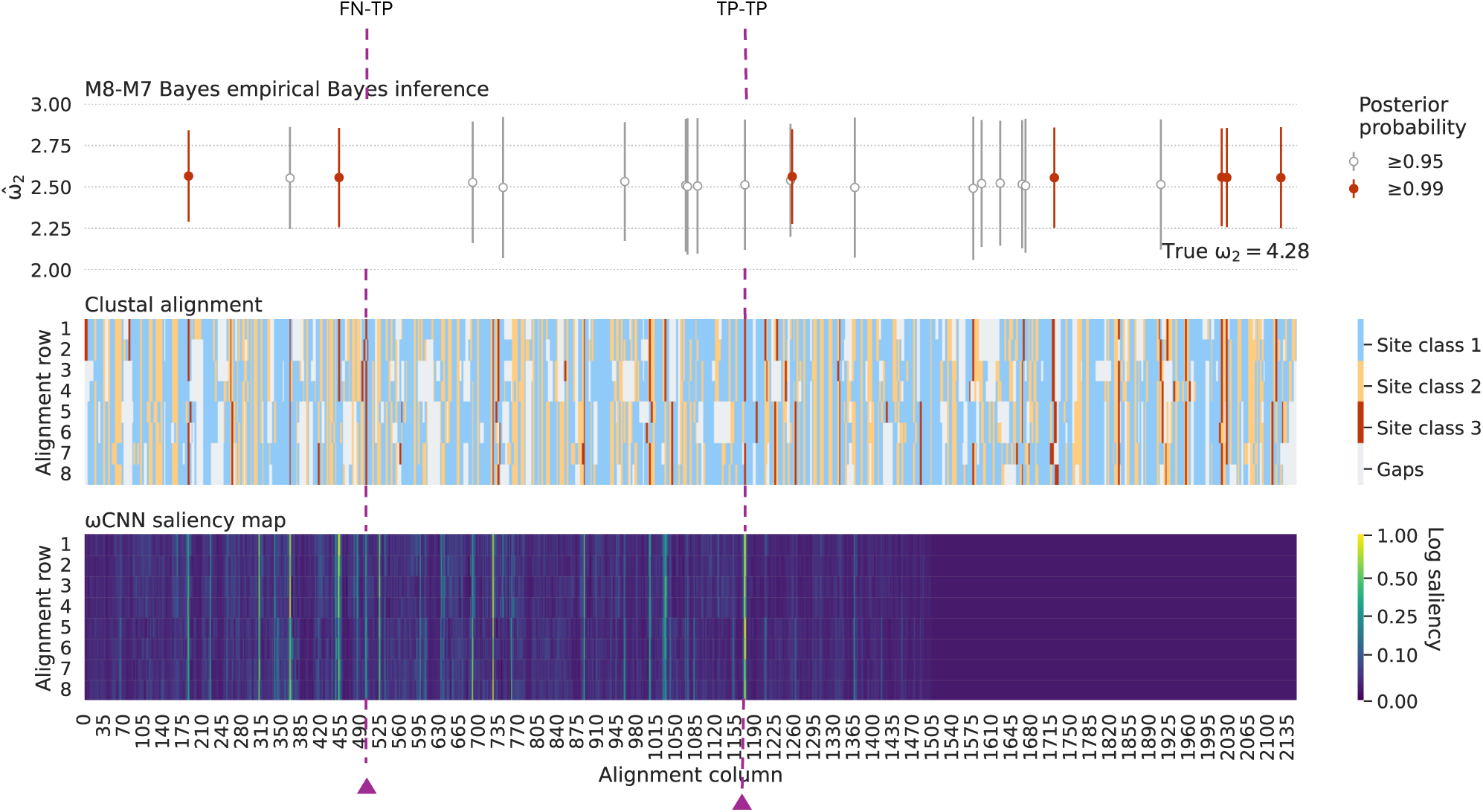
Saliency map for an MSA of protein-coding sequences evolving under positive selection, compared with the MSA and CODEML sitewise predictions of positive selection. Top panel: CODEML sitewise predictions of positive selection. Following significant results from both M1a/M2a and M7/M8 LRTs (as described in Methods), a Bayes empirical Bayes approach is used to calculate the posterior probability that each site is from a particular site class. Inferences of sites belonging to site classes with *ω >* 1 with *p* > 0.95 or *p* > 0.99 are shown in grey and red bars, respectively. **Middle panel:** Colour coded Clustal alignment. Blue, yellow and red sites indicate the three site classes described in the Methods, with red indicating sites with *ω >* 1. Grey indicates gaps in the alignment. **Bottom panel:** Saliency map, with bright regions (high saliency) showing the regions of the MSA that are most influential for OmegaAI’s classification. **Vertical lines:** The vertical line labelled TP-TP is an example alignment location of both CODEML and OmegaAI correctly identifying a site undergoing positive selection. FN-TP is an example of CODEML failing to detect a site undergoing positive selection whereas OmegaAI correctly detects positive selection at this location. Further examples are highlighted in Supplementary Fig. 11.

The correlation between sites under positive selection and high saliency can be leveraged as a proxy for sitewise detection of positive selection by OmegaAI, analogous to the BEB approach used by CODEML. Assuming this proxy, we can contrast the differences in sitewise predictions between CODEML and OmegaAI. Whilst the two methods tend to agree overall, it is informative to look at some of the details. In Figure 9 we highlight two examples: firstly, a site (labelled TP-TP) where CODEML and OmegaAI both correctly infer a site with *ω* > 1, despite some misalignment in the area; and secondly, a site (labelled FN-TP) where the OmegaAI saliency correctly suggests positive selection that CODEML fails to detect. Supplementary Fig. 11 provides an extended analysis of classification instances where CODEML and OmegaAI agree and disagree for this saliency map. We note the low saliency in the far right of the MSA and postulate that this is a result of zero-padding during training (see Discussion).

### 3.7 Computational resource measurements

A drawback of likelihood-based methods is the computational time required for parameter estimation. This becomes increasingly significant when scaling up the analysis to many genes or trees. Here we show that machine learning approaches can offer an alternative that is time and compute efficient.

In Figure 10, we present the computational time and memory usage for evaluating a standard test set of 2,000 alignments using CODEML and OmegaAI with baseline parameters. OmegaAI is evaluated in two scenarios: one using only CPU resources, and another utilising a single GPU in addition to a CPU.

**Fig. 10.**
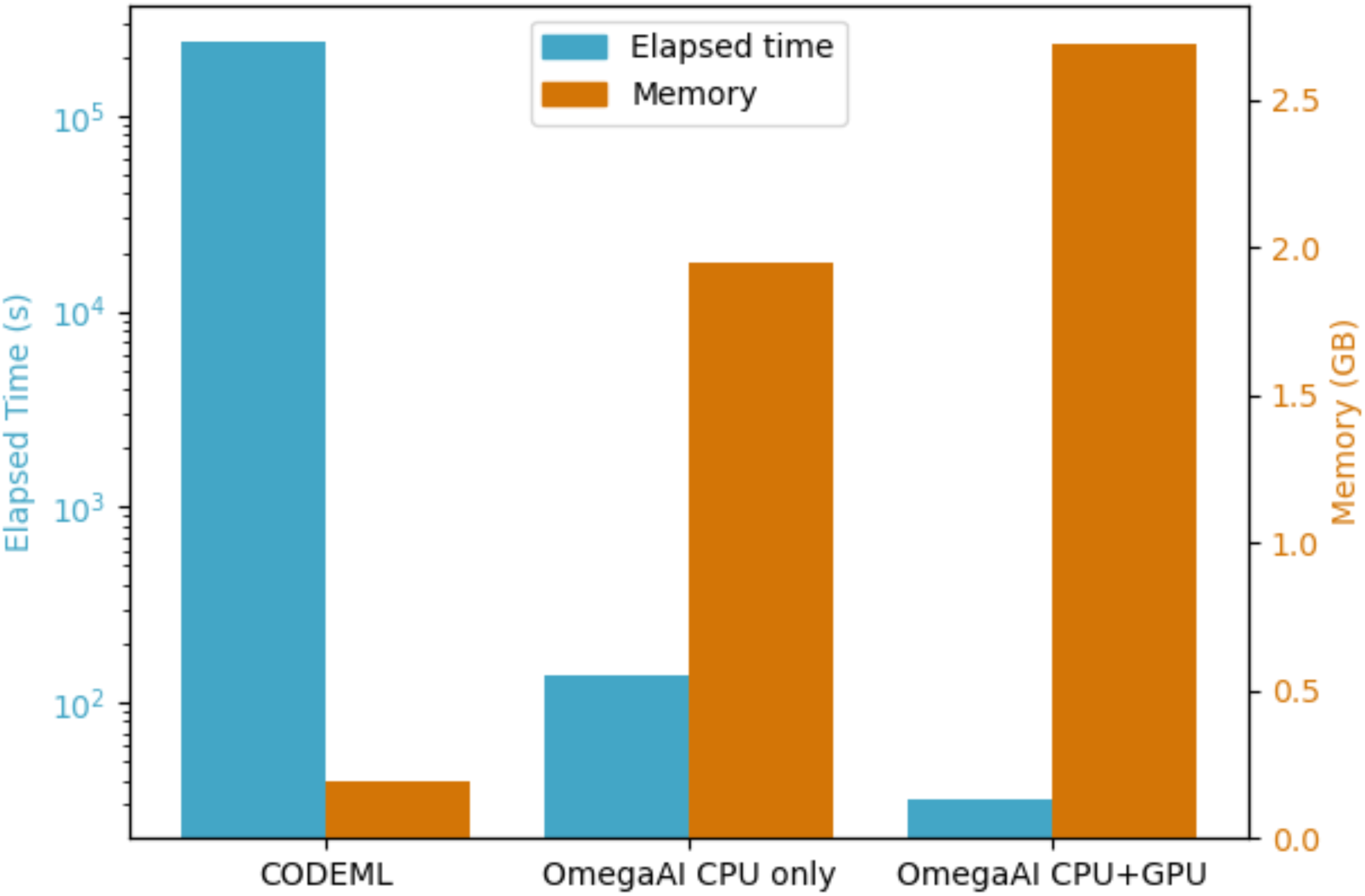
OmegaAI vs CODEML resource usage. These results are from 2,000 test MSAs (simulated under baseline parameters) sequentially evaluated by OmegaAI and CODEML. OmegaAI results are given for two scenarios: one where there is only a single CPU available and the other where both a CPU and GPU are utilised. OmegaAI requires the test data to be in TFrecord format (see Methods) before analysis and measurements presented here exclude the minimal resources required for pre-processing into this condensed format.

OmegaAI exhibits better speed, being four orders of magnitude faster with an average processing time of approximately 32 seconds to evaluate the 2,000 alignments when given access to the parallel processing capability of a GPU. In contrast, CODEML takes around 65 hours to evaluate these alignments when executed serially on a single CPU. When OmegaAI is restricted to a single CPU its processing time increases somewhat, to approximately 137 seconds in this example.

While the resource demands during training OmegaAI models (Supplementary Fig. 12) are substantial, the advantage of OmegaAI during test time is evident. This efficiency comes with a trade-off: OmegaAI requires more memory than CODEML, although the amount remains well within the capacity of modern personal computers. However, since OmegaAI lacks generalisability over tree shapes and divergences and would require retraining or at least fine-tuning for specific use-case utility, OmegaAI does not yet realise large advantages over CODEML in terms of resources. A possible exception to this is genome-scale detection of positive selection (see Discussion).

## 4 Discussion

The OmegaAI CNN, trained to perform positive selection detection on a set of aligned, interspecific, protein-coding nucleotide sequences, appears to outperform the state-of-the-art maximum likelihood methods, as embodied in the CODEML program in the PAML package (Yang, 2007), when trained for a specific phylogenetic scenario. This is particularly clear when dealing with high divergences and indel rates.

Using Clustal alignments to train OmegaAI is time efficient and, because Clustal alignments are more error-prone, potentially provides a richer resource from which the CNN model can learn to account for alignment error. Once an OmegaAI model has been trained, it provides a significant time advantage for dataset evaluation compared to maximum likelihood methods. These are promising results, suggesting that machine learning approaches could be valuable for detecting selection in multiple sequence alignment datasets. It is noteworthy that using a quicker, albeit less accurate, aligner during training for OmegaAI does not seem to present significant drawbacks. In fact, it substantially reduces training data generation time and the resulting OmegaAI model exhibits good performance even when evaluating lower-quality alignments, without a considerable drop in performance with higher-quality alignments. Where OmegaAI has been trained on MSAs containing alignment errors, we postulate that it has been able to learn patterns associated with positive selection, including selection signals made noisy by alignment error, effectively allowing OmegaAI to learn to accommodate alignment errors.

Although OmegaAI is not yet generalised to deal with more varied scenarios, particularly with respect to the number and evolutionary relationships of input sequences, this work serves as a proof of concept. Through a deliberately constrained version of the positive selection inference problem, we have begun to explore the capabilities of AI methods applied to this task. At present, the efficacy of OmegaAI relies on retraining for parameters specific to a given problem. For example, a feature already available in CODEML is that it can estimate divergence level during selection detection. Our results suggest that for OmegaAI it may be necessary to train new models using data simulated under a divergence level close to that of the target data, which in of itself requires approximate prior knowledge of the divergence level of the test data, or using more extensive data drawn from a range of divergence levels. Another mitigating strategy could be to fine-tune existing models instead of fully retraining, which appears to be a promising strategy based on the results from the mixed divergence OmegaAI model.

The prospects for creating a generalised AI model capable of high performance over many different problems look promising and could conceivably be aided by an attention-based architecture or by increasing the size of the model. Attention-based architectures have proven to be promising in a range of tasks, including applications to MSAs (Rao et al., 2021; Jumper et al., 2021). Larger machine learning models — those with more parameters and deeper networks — tend to perform better in terms of generalisation because they have more capacity to learn complex patterns and representations from data, allowing them to better capture the underlying distribution and make accurate predictions on new, unseen examples.

One use-case for which OmegaAI suggests immediate utility may be genome-wide detection of positive selection. In these investigations, typically thousands of genes are tested for positive selection. For example, Lee et al. (2017) query over 1,000 genes in nine primates using likelihood-based methods. The underlying species tree is constant and mammalian gene evolutionary rates tend to be approximately normally distributed with a similar range to the divergence levels that we tested (Kumar and Subramanian, 2002), and so a small number of OmegaAI models could be trained to cover this range. At this scale, even including the time it takes to simulate training data and train the model(s), OmegaAI could present a time-efficient solution that is potentially able to provide more accurate inferences, according to our results.

Our results show that OmegaAI retains some predictive power with difficult datasets, where the likelihood methods that we have tested perform poorly. In particular, it shows promise for noisy data that result from high divergence and indel rates. This too is an encouraging potential use-case for OmegaAI, where current statistical models are likely to be of little use because of their failure to account for alignment errors.

Another interesting avenue to investigate would be alternative evolutionary models for simulating data. The simulation model should reflect biological reality in order for OmegaAI models to perform well on empirical data. The space of models available for simulating sequence evolution is larger and more complex than those used for maximum likelihood optimisation (McGuire, Denham, and Balding, 2001) and increasingly complex models can better capture evolutionary mechanisms and emergent properties (De Maio et al., 2013; Perron et al., 2019; Lucaci et al., 2021). Since our AI model is trained on simulated sequences, this allows for training on sequences generated under complex and realistic models, even those intractable for likelihood inference. This could lead to a machine learning model that makes more accurate inferences on empirical data. For example, we could simulate the phenomenon of multiple nucleotide substitutions (knowledge of which has been shown to reduce the rate of false inference of positive selection: De Maio et al., 2013), take into account the growing evidence for the non-neutrality of synonymous substitutions (Parmley, Chamary, and Hurst, 2006; Kubatko et al., 2016; Shen et al., 2022; Rosenberg, Marx, and Bronstein, 2022), and model synonymous rate variation (Wisotsky et al., 2020) and interlocus gene conversion (Ji, Griffing, and Thorne, 2016). Machine learning approaches are often preferable to approximate Bayesian computing (ABC) — an alternative to likelihood-based estimation for intractable models — because they automate the data-to-parameter mapping, eliminating the need for selecting summary statistics and comparison metrics. Additionally, machine learning methods are generally more scalable, particularly for large datasets (Lenzi et al., 2023).

Finally, it is a natural extension to imagine a machine learning model whereby we try to infer sitewise d N/d S, a feature already available in the likelihood framework, for example as implemented within CODEML through empirical Bayes approaches or in SLR (Massingham and Goldman, 2005) through maximum likelihood. In the CNN context, this problem is akin to a pixel-classification problem already attempted in Längkvist et al. (2016), or could be approached using an attention-based architecture, similar to Nesterenko et al. (2024). To simultaneously achieve sitewise detection of positive selection and sequence length generalisability, however, our saliency maps have uncovered that extra care is needed beyond crude zero-padding (see Methods). We regularly observe weaker saliency towards the far right end of MSAs, likely attributable to zero-padding that was necessary to accommodate variable lengths in sequences during training. The CNN, regularly encountering zeros in these positions across numerous training datasets, appears to have learned to prioritise areas most frequently containing relevant information. There is enough signal in higher saliency regions that binary classification performance has not been strongly impacted, but in sitewise inference we would require uniformly high signal across the whole length of an MSA. A sliding window approach, although potentially less efficient, could be used to address this problem by querying sections of the MSA with some overlap to capture spreading of horizontal information from misalignment.

In conclusion, machine learning methods applied to detecting interspecific positive selection have demonstrated superior accuracy and time efficiency compared to likelihood methods, when trained appropriately for a specific problem. In this study, we concentrated on training and testing models for particular use-cases, observing some limitations in generalisation outside of the trained scenarios. The application of current and future machine learning techniques, which are more sophisticated, is expected to enhance generalisability and thus increase their utility for a broader range of phylogenetic topologies and questions. Although machine learning methods still face “black box” issues, we have begun to address this through saliency map exploration. Whilst likelihood methods remain valuable due to their strong statistical foundations and extensive usage, the adoption of learning-based methods is likely to increase due to their growing transparency, scope, and sophistication.

## 5 Methods

### 5.1 Simulating multiple sequence alignments

Following earlier studies which investigated the effects of insertions, deletions, and alignment errors on tests of positive selection (Fletcher and Yang, 2010; Jordan and Goldman, 2012), we simulate codon sequences under artificial trees for a range of testing scenarios using INDELible (Fletcher and Yang, 2009). This tool, given a variety of user-defined parameters, outputs both unaligned sequences and the true alignment of those sequences. Alignments of these simulated sequences act as our training, validation, and test datasets for our CNN and as our test sets for evaluation using the likelihood-based methods for detecting selection implemented in CODEML from the Phylogenetic Analysis by Maximum Likelihood (PAML; Yang, 2007) package. Unlike traditional classification tasks in deep learning, which are limited by the amount of available training data, we are able to use simulations to generate as many pairs of alignments and true class labels (selection or no selection) as desired for each testing scenario.

Considering a constrained version of the problem, we used an 8-taxon symmetric topology rooted at its midpoint (similar to those used in Fletcher and Yang, 2010; Jordan and Goldman, 2012). The intention is to mimic the evolution of a protein-coding gene sequence along an idealised tree under varying selection pressures. Some parameters, and particularly sampling distribution parameters and ranges, are fixed across all simulations. The sequence length at the root of the tree is sampled from a gamma distribution with parameters k = 4.2 and *✓* = 85, which approximates bacterial gene length distributions as outlined in Skovgaard et al. (2001). To prevent sampling of extreme lengths, we impose minimum and maximum root sequence lengths of 100 and 600 codons, respectively. Additionally, the ratio of transitions to transversions, *K*, is sampled from U(2, 3), the continuous uniform distribution with a minumum value of 2 and a maximum value of 3. This range is close to that which is observed in empirical studies (Wang et al., 2015).

Sequences were simulated using a version of the codon model of evolution with site variation in selective pressure from Nielsen and Yang (1998). The model assumes that each codon alignment position is a member of a discrete site class S. Each site class is assigned a different level of selective pressure as measured by the nonsynonymous to synonymous substitution ratio, *ω*, with selective pressure being constant across all branches of the tree at a considered codon position. We assume three site classes S 2 S_0_, S_1_, S_2_ occurring in proportions p_0_, p_1_, p_2_, respectively, each associated with different *ω* values. The value of *ω*_0_ for S_0_ is chosen from U(0.1, 0.5), and *ω*_1_ for S_1_ is chosen from U(0.5, 0.9). Sites in S_0_ are under strong purifying selection and those in S_1_ are under slight purifying selection. The value of *ω*_2_ for S_2_ is chosen differently depending on the type of gene we are simulating; genes that are predominantly undergoing purifying selection (*ω*_2_ 2 U(0.9, 1.0)), genes containing strictly neutral sites (*ω*_2_ = 1), and genes where some sites are evolving under positive selection (*ω*_2_ 2 U(1.5, 5)). For simplicity, in this study we refer to these three types of simulated genes as purifying, neutral and positive selection genes, respectively. The model used to simulate genes under purifying and neutral evolution is similar to the models M1a and M7 implemented in CODEML, while the model used to simulate genes under positive selection is similar to M2a and M8 (Nielsen and Yang, 1998; Yang, Nielsen, et al., 2000; Yang, Wong, and Nielsen, 2005).

For every gene, we sample one value each for *ω*_0_, *ω*_1_, and *ω*_2_ from their corresponding distributions. This ensures that for each gene, every codon site will evolve under one of three *ω* values and sites of the same site class will experience identical selection pressure. For clarity, the distributions of *ω* are shown for our baseline parameters for each type of selection in Supplementary Fig. 13. The proportion of class S_0_ sites, p_0_, was chosen independently for each gene from U(0.5, 0.8), and p_2_ was sampled from U(0.01, 0.1); p_1_ is then set as p_1_ =1 - p_0_ - p_2_.

The branch lengths of the ultrametric simulation tree are all equal, varying between 0.1 and 1.0 in different simulations, with a baseline value of 0.2 (see Supplementary Fig. 1). These branch lengths represent the expected number of substitutions per codon site per branch. The ratio of insertions to deletions was fixed at 1, and the rate of insertions and deletions (indels) to substitutions was set at 0.1 (close to the empirically determined value for coding regions in mammals; Chen et al., 2009) as part of the baseline parameter set. More extreme values of 0.2 and 0.3 were also tested (rates closer to those observed in bacteria: Chen et al., 2009), with results shown in Supplementary Fig. 6 and Supplementary Fig. 7, respectively. As in Fletcher and Yang (2010), indel lengths are drawn from a geometric distribution with parameter q = 1 - p = 0.35, as this provides a good fit to observed indel length distributions in protein coding sequences (Taylor, Ponting, and Copley, 2004).

Using the procedure outlined above, for each unique set of parameter values, we simulated 1,000,000 alignments for training and validating our network, and an additional 2,000 alignments to test and compare the performance of our network and the likelihood-based methods provided by CODEML (Yang, 2007). For training and testing simulations, 40% of the genes for each set of parameter values are evolved under negative selection, 10% with neutral selection, and 50% with positive selection. These proportions are chosen to achieve a balanced training dataset and avoid bias towards a majority class (Wei and Dunbrack, 2013; Ghosh et al., 2024).

All generated nucleotide sequences are converted into their corresponding amino acid sequences before further processing in order to maintain reading frame between sequences. The training and validation sets are aligned using Clustal Omega v1.2.4 (Sievers, Wilm, et al., 2011), which is fast, but is known to be capable of producing a high false positive rate when inferring selection using traditional methods (Löytynoja and Goldman, 2008; Fletcher and Yang, 2010). For each of the 2,000 testing alignments, we retain the true alignment, re-align each set of sequences using Clustal Omega, and additionally align using MAFFT v7.475 (Katoh and Standley, 2013) and two variants of PRANK v170427 (Löytynoja and Goldman, 2008), one using an amino acid model (subsequently referred to as PRANKaa) and one using an empirical codon model (subsequently referred to as PRANKc). All amino acid alignments are converted back into their original nucleotide sequences using PAL2NAL v14 (Suyama, Torrents, and Bork, 2006).

### 5.2 Alignment transformation

Multiple sequence alignments are typically represented as an m ⇥ n matrix, of m sequences and n alignment columns, with each alignment position in this matrix occupied by a nucleotide or gap character. It is necessary to encode this information numerically before passing it as input to any CNN. Ideally, encoding should avoid introducing any numerical relationships between entries in the input data which were not present before encoding; for example, encoding nucleotides with integers 0, 1, 2, and 3 would artificially introduce different distances between different pairs of nucleotides. To avoid this, we use one-hot encoding of our input alignment matrix, where each nucleotide/gap character is converted into one of five unique bit patterns (A : 00001, C : 00010, G : 00100, T : 01000, - : 10000). We then convert our m ⇥ n matrices (where m = 8 and n =3 ⇥ number of codons in the alignment) into three-dimensional m ⇥ n ⇥ 5 tensors recognised by the CNN, where the third dimension of each alignment position contains the five bits just described, corresponding to the character present at that position. (This is analogous to the common 3D encoding of RGB images in which the third dimension represents composite colour via three distinct colour channels.) For training, we store these encoded alignments (along with their true class label) in TFRecord format, a binary file format used for efficient serialisation of structured data (Abadi et al., 2016).

### 5.3 OmegaAI: architecture

Here we describe our final network structure, which we refer to as OmegaAI. Before arriving at our final architecture and hyperparameter selection we conducted preliminary tests using 100,000 Clustal Omega alignments, simulated under baseline conditions. We selected the architecture with the highest F_0.5_ score on this test set, which was an architecture with six convolutional layers. The F_0.5_ score provides a measures of accuracy for binary classification tasks that places a greater weight on precision rather than recall. The learning rate is a hyperparameter that determines the size of the steps the model takes to adjust its weights during training, affecting the speed and stability of the learning process. A learning rate of 0.001 was opted for as the baseline, having tested a standard range. See Supplementary Fig. 14 for details of these preliminary tests.

All network implementation was done in Python v3.6.12, using Keras v2.3.0, which is packaged with TensorFlow v2.2.0 (Abadi et al., 2016). Our network is structured similarly to that of Suvorov, Hochuli, and Schrider (2020), but uses six convolutional layers instead of eight. The first layer uses a filter size of m ⇥ 3, where m is the number of sequence rows (taxa) in each dataset, and 3 corresponds to the three nucleotides of each codon alignment column. We use a stride of 3 in the first layer to evaluate each codon column separately. We tested using a smaller filter size of 3⇥3 for the first layer, but found that convolutions over all m sequence rows resulted in better performance. We also tested a smaller stride size to allow neighbouring partial codons to be evaluated (i.e. still with filter size m ⇥ 3, but now the convolution can also covers 1 or 2 columns from one codon, and correspondingly 2 or 1 from an adjacent codon), but noticed no appreciable improvement in performance. Note also that operating over each m ⇥ 3 column with no overlap permits faster training than using smaller values for filter size and stride.

After the convolutional layers, we use global average pooling across the third dimension of the weight tensors. We chose this pooling operation over the commonly used flatten layer, as it reduces the spatial dependency between specific input alignment regions and the resulting classifications (Lin, Chen, and Yan, 2013). As our problem is one of binary classification (selection, 1 vs. no selection, 0), we use a sigmoid activation function in our final dense layer. Other than for our initial layer, in which convolutions are over codon columns (see above), filter sizes follow those outlined in Suvorov, Hochuli, and Schrider (2020). After each convolutional layer, we use a standard batch normalisation (Ioffe and Szegedy, 2015), dropout regularisation (p = 0.5; Srivastava et al., 2014), then average pooling. This relatively high dropout probability of 0.5 was used because lower rates tended to produce overfitted networks which performed worse when evaluating test datasets (Supplementary Fig. 14).

Our aim was to enhance accuracy and reduce false positives in Clustal Omega alignments simultaneously. Each model underwent 50 epochs of training, with evaluation conducted on the model at the 50th epoch. Stopping at 50 epochs means we stop before validation accuracy falls due to over-fitting (see Supplementary Fig. 15).

### 5.4 OmegaAI: training

All training runs were performed on a single GPU (NVIDIA Tesla V100 PCIe 32 GB), within a highperformance computing environment. Weights in our network are initialised using Glorot uniform initialization (Glorot and Bengio, 2010). As is typical when working with large training datasets (here, storing 950,000 training alignments in memory is almost always unfeasible), we use nonoverlapping, subsets or “batches” of alignments for training. Each of these batches consists of 512 randomly chosen alignments, zero-padded to the length of the longest alignment in that batch, as batched inputs to CNNs are required to have equal dimensions. For each batch, weight updates are performed using the Adam optimisation algorithm (Kingma and Ba, 2017) to minimise a binary cross-entropy loss function that distinguishes positive selection (1) from no selection (0). This procedure is performed for a total of 50 epochs, where one epoch has passed when all 950,000 training alignments have been seen once. Note that the alignments in each batch change at each epoch to avoid overfitting on any particular batch. Adam optimisation is well-established and generally performs well across a range of learning tasks (Soydaner, 2020). We also tried training using AdamW (Loshchilov and Hutter, 2017), a popular extension of Adam, but noticed no appreciable improvement in performance.

For the standard baseline OmegaAI model, solely Clustal alignments are used to train the CNN. Clustal is fast (Sievers, Wilm, et al., 2011) which is crucial for the scalability needed in generating large amounts of training data. In addition, the tool is also widely used and accessible. Despite being the most error-prone among the aligners tested (see Supplementary Fig. 2 and Supplementary Fig. 3), this characteristic presents an opportunity for the CNN’s training process and may even be beneficial (see Results and Discussion).

### 5.5 Comparison of OmegaAI with maximum likelihood methods

To evaluate the performance of both OmegaAI and likelihood-based tests for selection, we use the true, Clustal Omega, MAFFT, PRANKc, and PRANKaa alignments generated for each parameter set (2,000 alignments for each aligner).

In all tests, we perform classification under OmegaAI on each of the 2,000 Clustal Omega alignments, and in most cases, we also classify the alignments of the same sequences obtained from all other aligners. For almost all test sets, and consistent with our network training, we apply a threshold of 0.5: the classification of an alignment is ‘positive selection’ if the output value from the final sigmoid activation layer, Z, satisfies Z > 0.5, and ‘no selection’ otherwise. For a subset of test sets, we also explore increasing the threshold for calling positive selection in order to reduce the false positive rate.

Likelihood-based tests are facilitated by the CODEML program implemented in PAML (Yang, 2007), which provides statistical methods for detecting the presence of positive selection in alignments of protein-coding nucleotide sequences under a variety of statistical models of selection. We used the two recommended pairs of models for detecting selection in scenarios acknowledging the possible presence of sitewise variation in *ω*: M1a/M2a (Nielsen and Yang, 1998; Yang, Wong, and Nielsen, 2005) and M7/M8 (Yang, Nielsen, et al., 2000). Each pair enables a likelihood ratio test (LRT) between hypotheses of all sites evolving with no positive selection (*ω* 6 1), and evolution with positive selection (*ω* > 1 at some sites). Briefly, in the first LRT, M1a assumes two site classes, *ω*_0_ < 1 and *ω*_1_ = 1, while M2a assumes three site classes, *ω*_0_ < 1, *ω*_1_ = 1, and *ω*_2_ > 1. Note the correspondence between the assumptions of M1a and M2a and the distributions shown in Supplementary Fig. 13. In the second LRT, M7 assumes that all sites are conserved with *ω* ⇠ beta(p, q), while M8 assumes that a proportion (p_0_) of sites are conserved with *ω*_0_ ⇠ beta(p, q), and all other sites (p_1_ = 1 - p_0_) evolved under positive selection with *ω* > 1. The parameters in all of these models are estimated by maximising their likelihood functions (see Yang, Nielsen, et al., 2000), and the significance of the LRT between no selection and selection is evaluated using a *x*^2^ test with two degrees of freedom in each case. In this study, a significance level of 0.05 is chosen, indicating a 95% confidence level for determining the presence of positive selection, which is a common threshold in the literature (Wong, Yang, et al., 2004). We define CODEML’s classification of positive selection to be the result where both the M1a/M2a and M7/M8 LRTs produce significant, positive results. We do not classify a gene to be under positive selection if one or both LRTs are non-significant. CODEML is given the simulation topology as a start tree and is allowed to re-estimate branch lengths during the likelihood maximising process: CODEML tends to perform better under this regime compared to enforcing fixed branch lengths (see Supplementary Fig. 16).

In addition to enabling LRTs for binary classification of a whole MSA, CODEML can conduct a Bayes empirical Bayes analysis to estimate sitewise *ω* values in cases where we want to study the specific location(s) of signal for selection within alignments (Yang, 2007; Álvarez-Carretero, Kapli, and Yang, 2023). We use this for comparison with the saliency approach for CNNs, described in the next section.

### 5.6 Alignment saliency

We are interested in where the most informative positions lie within an MSA input, and in particular what the characteristics of these informative positions are. We can investigate this by quantifying the importance of each position in an input alignment on the prediction from our network using saliency maps (Simonyan, Vedaldi, and Zisserman, 2014). For example, we can expect that the most informative sites for predicting positive selection are those evolving with *ω* > 1. We can also examine the saliency of misaligned and neighboring columns, particularly where codon sites that have evolved under different selection pressures are incorrectly placed in the same column. Given a CNN producing an output score Z (between 0 and 1, ultimately thresholded to infer selection or no selection) from any input alignment A (of size m sequences ⇥ n alignment columns ⇥ 5 nucleotide/gap characters), a saliency map is a function M which assigns an importance M(A)*_i,j_* 2 R to each alignment position (i, j) based on the influence of that position on the output score Z(A). We use the saliency function of Simonyan, Vedaldi, and Zisserman (2014) as it provides a suitable method for calculating the sitewise saliency for one-hot encoded values. Given a specific alignment A_0_ and associated binary score Z(A_0_), the (non-linear) function Z(A) is approximated as a linear function in the neighbourhood of A_0_ using the first-order Taylor expansion:

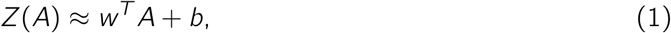

where w is the derivative of Z with respect to alignment A at A_0_:

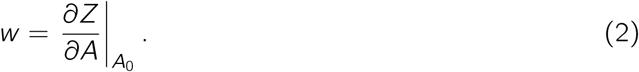

Note that in our application this derivative is three-dimensional (w = w*_ijc_*), because A has dimensions m ⇥ n ⇥ 5. To compute the full saliency map (M 2 R*^m^*^⇥^*^n^*) for each alignment, we first calculate the derivative w (Equation 2) by back-propagation, and then take the maximum magnitude of w across all nucleotide channels c at each alignment position (i, j) using

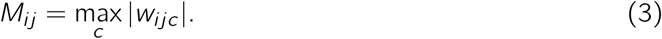

## Acknowledgments

We thank José Almeida for assistance with technical details of CNN architecture design and interpretation of results.

