## Supplementary Information for "Detecting interspecific positive selection using convolutional neural networks"

### 6 Supplementary Information

#### 6.1 Supplementary Figures

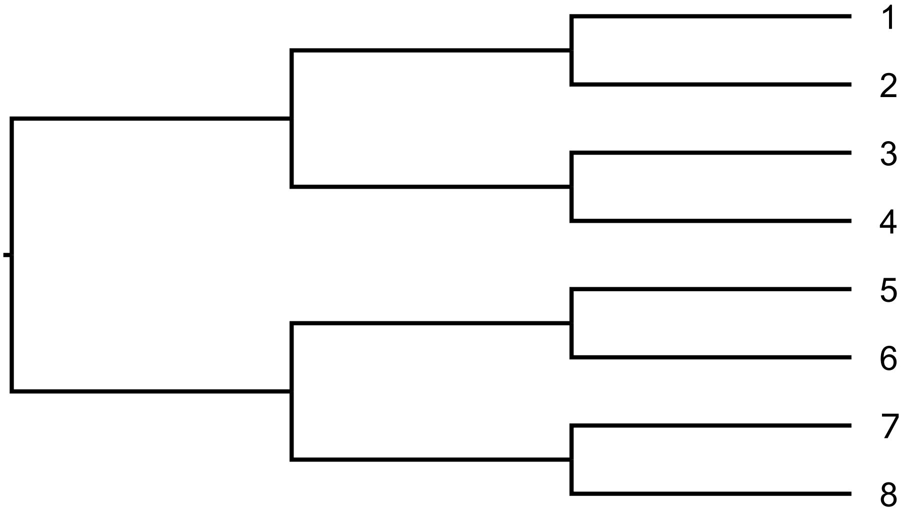


**Figure S 1. Artificial phylogenetic tree used for simulation.** An 8-taxon, symmetric, ultrametric tree with equal branch lengths scaled to a chosen divergence. Simulated sequences are evolved along this tree using INDELible (Fletcher and Yang, 2009) and associated MSAs have consistent row ordering relative to the ordering of the tips shown in the tree. For the baseline parameter set, each branch length is set to 0.2 expected substitutions per codon, making the root to tip length of the tree 0.6.

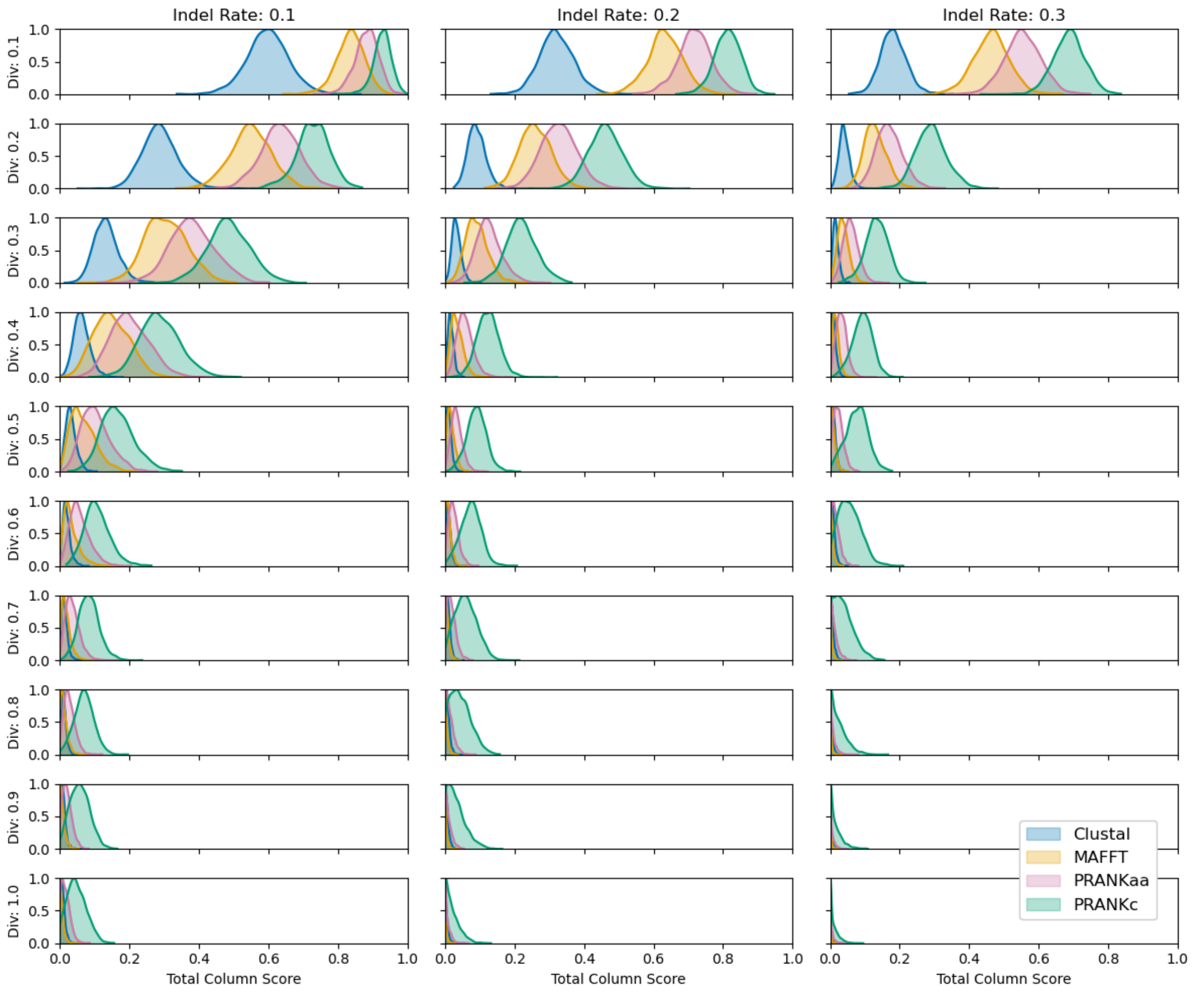


**Figure S 2. Alignment Total Column Scores across aligners, divergences and indels.** For each divergence-indel combination 2,000 sets of sequences along with their true MSAs were generated using INDELible (see Methods). For each set of sequences the alignment is then inferred using Clustal, MAFFT, and two PRANK versions (PRANKaa and PRANKc), forming the test sets in this study. The Total Column Score (proportion of true alignment columns in the inferred alignment) was calculated for each alignment using FastSP (Mirarab and Warnow, 2011) to assess alignment quality, which decreases as divergence and indel rates increase.

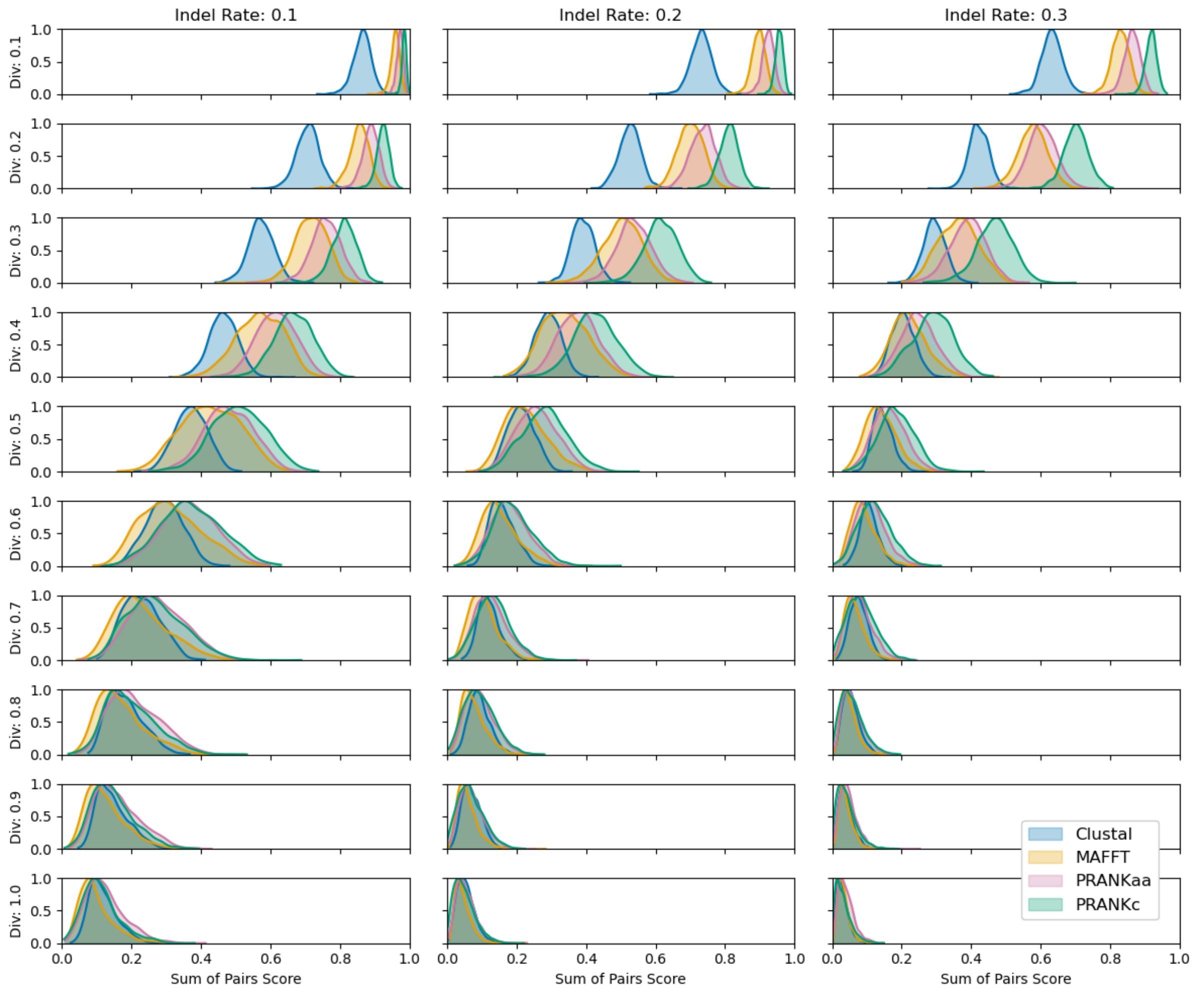


**Figure S 3. Alignment Sum of Pairs Scores across aligners, divergences and indels.** Details are as in Supplementary Fig. 2, except that here we show the Sum of Pairs Score, which is the proportion of true homologies (aligned pairs) found in an inferred alignment. The Sum of Pairs Score (here calculated with FastSP: Mirarab and Warnow, 2011), another measure of the quality of an alignment, decreases as divergence and indel rates increase.

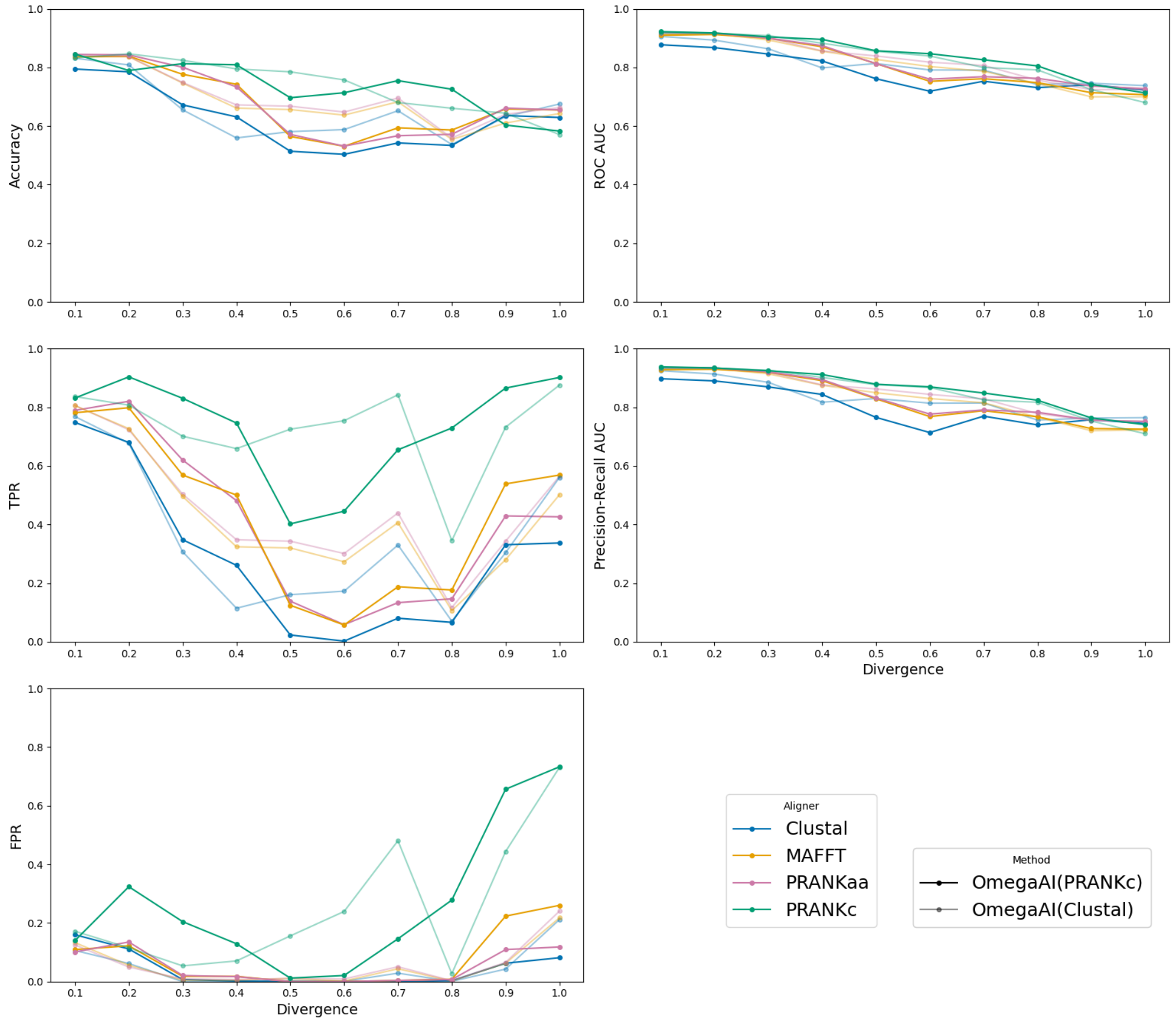


**Figure S 4. OmegaAI model trained on PRANKc alignments vs. OmegaAI model trained on Clustal alignments.** Various binary classifier performance metrics are presented to compare the two methods. In the first column of plots the accuracy, TPR and FPR are calculated under a threshold of 0.5 for both methods. The divergence axis refers to the scaling of branches of the 8-taxon symmetric tree (Supplementary Fig. 1) used for simulation, and a different OmegaAI model is trained for each divergence level and for each method. The baseline parameter for indel rate, 0.1, is used. The OmegaAI(Clustal) models are trained exclusively on Clustal alignments and shown in the semi-transparent lines, whereas OmegaAI(PRANKc) models are trained exclusively on PRANKc alignments and shown in the bold lines. The test data is aligned by the four different aligners and tested by both methods. The two sets of models perform comparably, indicating that there is little benefit from training on the more accurate PRANKc alignments, despite the considerable computational resource cost of running PRANK.

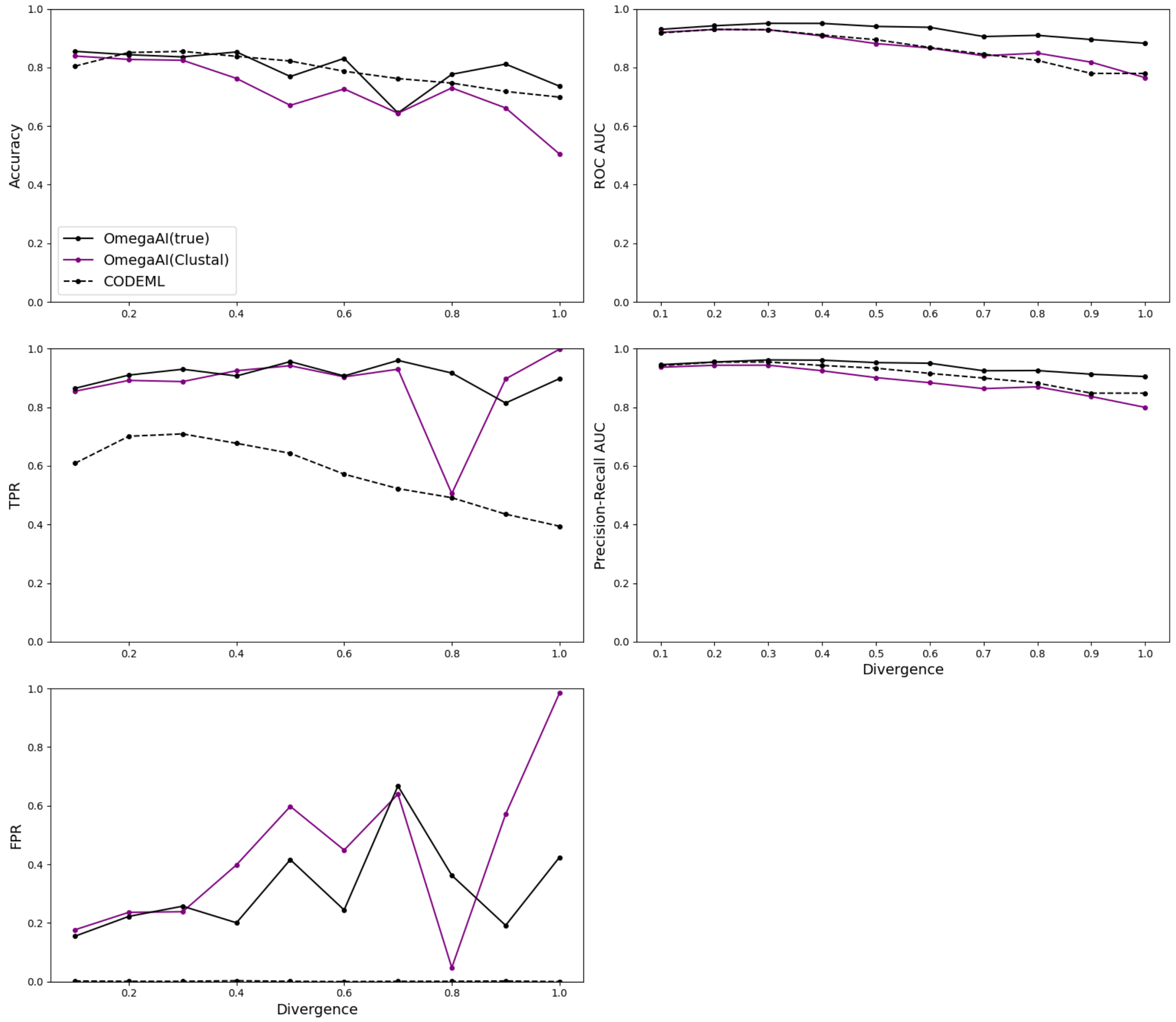


**Figure S 5. OmegaAI vs. CODEML — true alignments.** Various binary classifier performance metrics are presented to compare three methods: OmegaAI trained on true alignments or Clustal alignments, and CODEML. In the first column of plots the accuracy, TPR and FPR are calculated under a threshold of 0.5 for both AI methods, and a threshold of $p = 0.95$ for CODEML. The divergence axis refers to the scaling of branches of the 8-taxon symmetric tree (Supplementary Fig. 1) used for simulation, and a different OmegaAI model is trained for each divergence level and for each AI method. The baseline parameter for indel rate, 0.1, is used. All three methods are evaluated on the same set of 2,000 true alignments. In purple are the standard OmegaAI models trained on Clustal alignments. The bold black lines show results from OmegaAI models trained using true alignments; dashed lines are CODEML results. The three methods perform comparably, with the OmegaAI(true) models outperforming the standard OmegaAI models across metrics. The TPR and FPR suggest that the decision threshold could benefit from being lowered to reduce the FPR to come more in line with CODEML's conservative results.

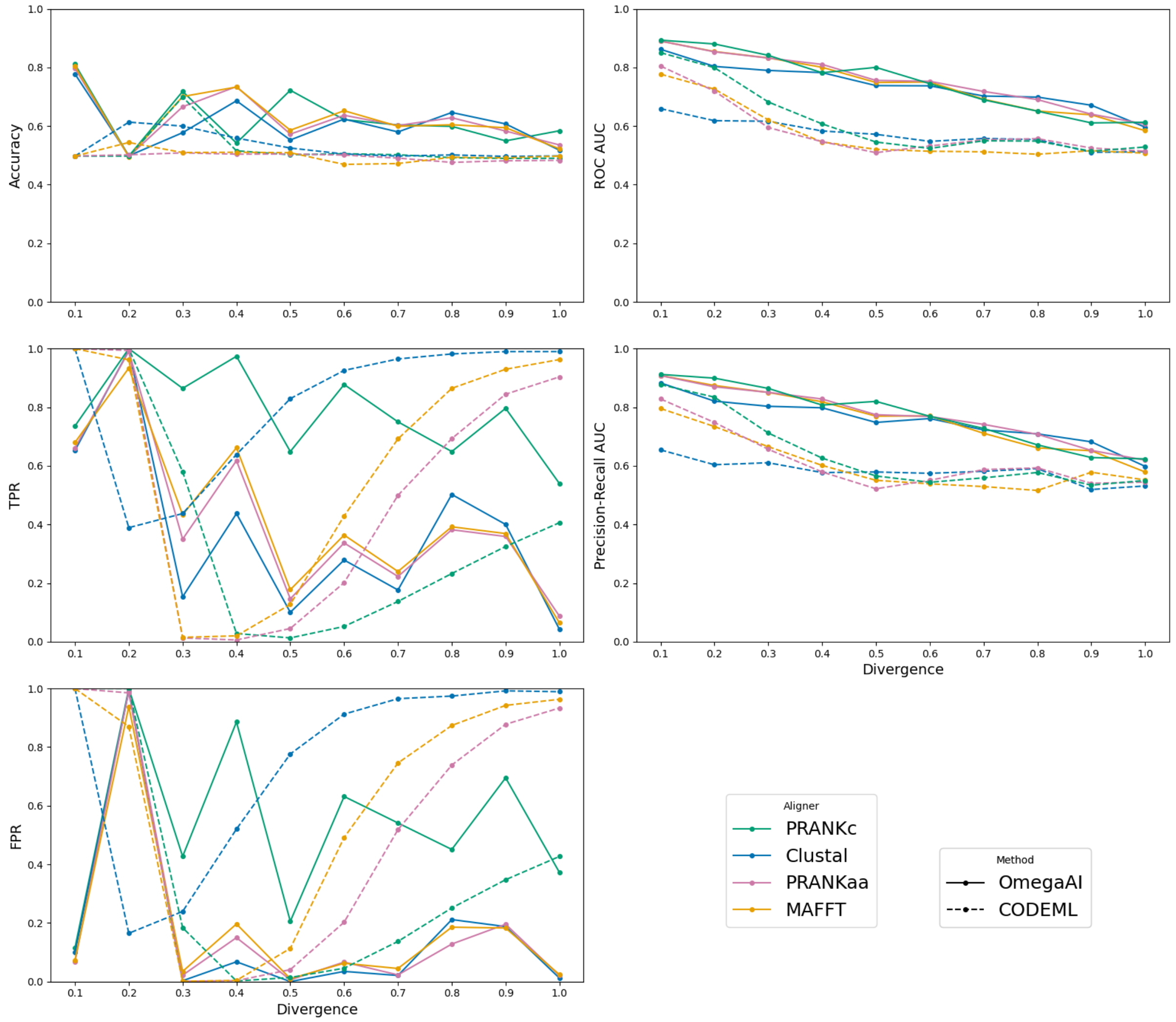


**Figure S 6. OmegaAI vs. CODEML — indel rate 0.2 across divergences.** Various binary classifier performance metrics are presented to compare the two methods. In the first column of plots the accuracy, TPR and FPR are calculated under a threshold of 0.5 for OmegaAI, and a threshold of $p = 0.95$ for CODEML. The divergence axis refers to the scaling of branches of the 8-taxon symmetric tree (Supplementary Fig. 1) used for simulation, and a different OmegaAI model is trained for each divergence level. A value of 0.2 is used for the indel rate. The OmegaAI models are trained exclusively on Clustal alignments. The test data is aligned by the four different aligners and tested by both methods. Increasing the indel rate from the baseline of 0.1 presents more variable trends and generally poorer performance from both methods.

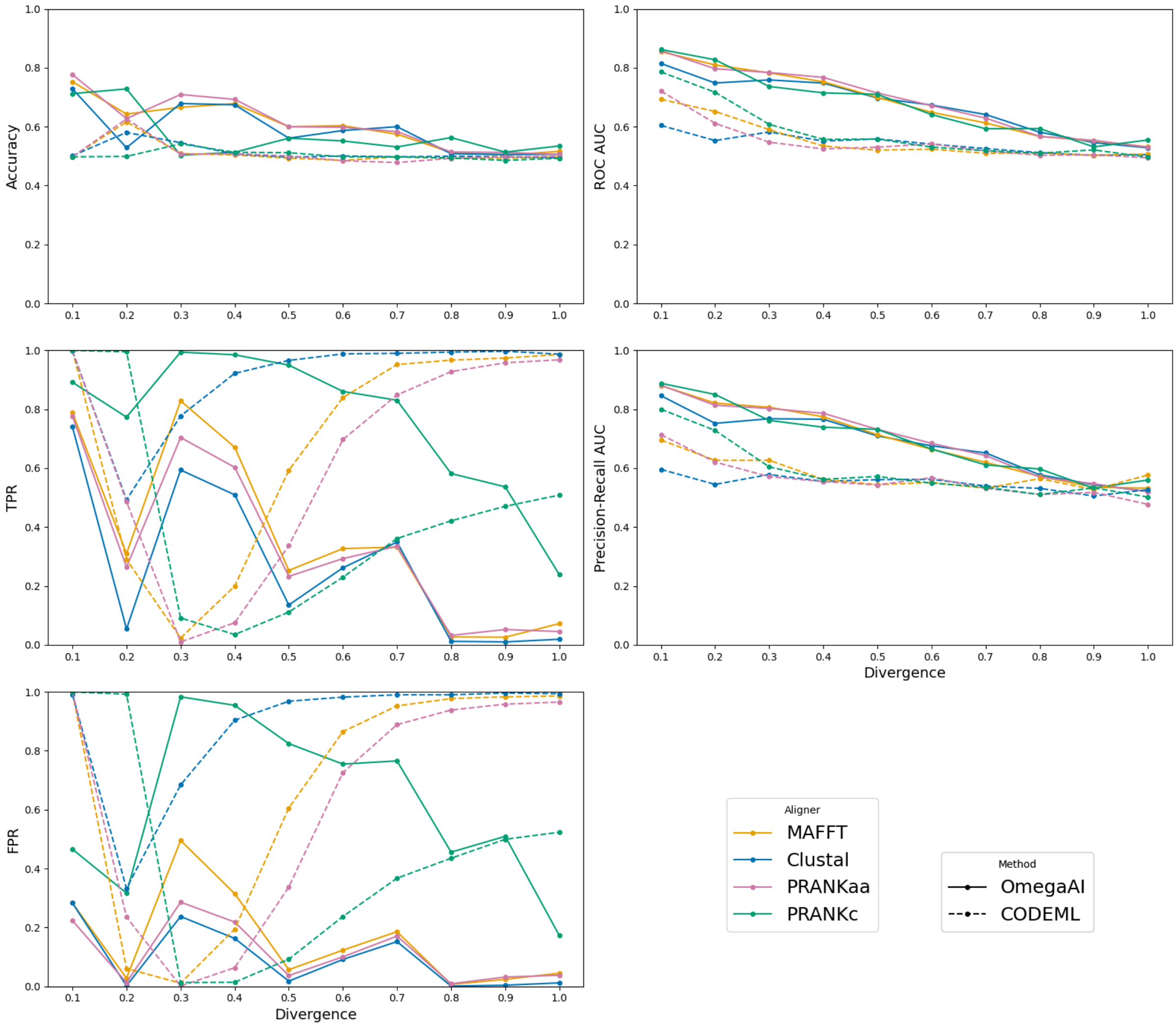


**Figure S 7. OmegaAI vs. CODEML — indel rate 0.3 across divergences.** Various binary classifier performance metrics are presented to compare the two methods. In the first column of plots the accuracy, TPR and FPR are calculated under a threshold of 0.5 for OmegaAI, and a threshold of $p = 0.95$ for CODEML. The divergence axis refers to the scaling of branches of the 8-taxa symmetric tree (Supplementary Fig. 1) used for simulation, and a model is trained for each divergence. A value of 0.3 is used for the indel rate. The OmegaAI models are trained exclusively on Clustal alignments. The test data is aligned by the four different aligners and tested by both methods. Increasing the indel rate from the baseline of 0.1 presents more variable trends and generally poorer performance from both methods.

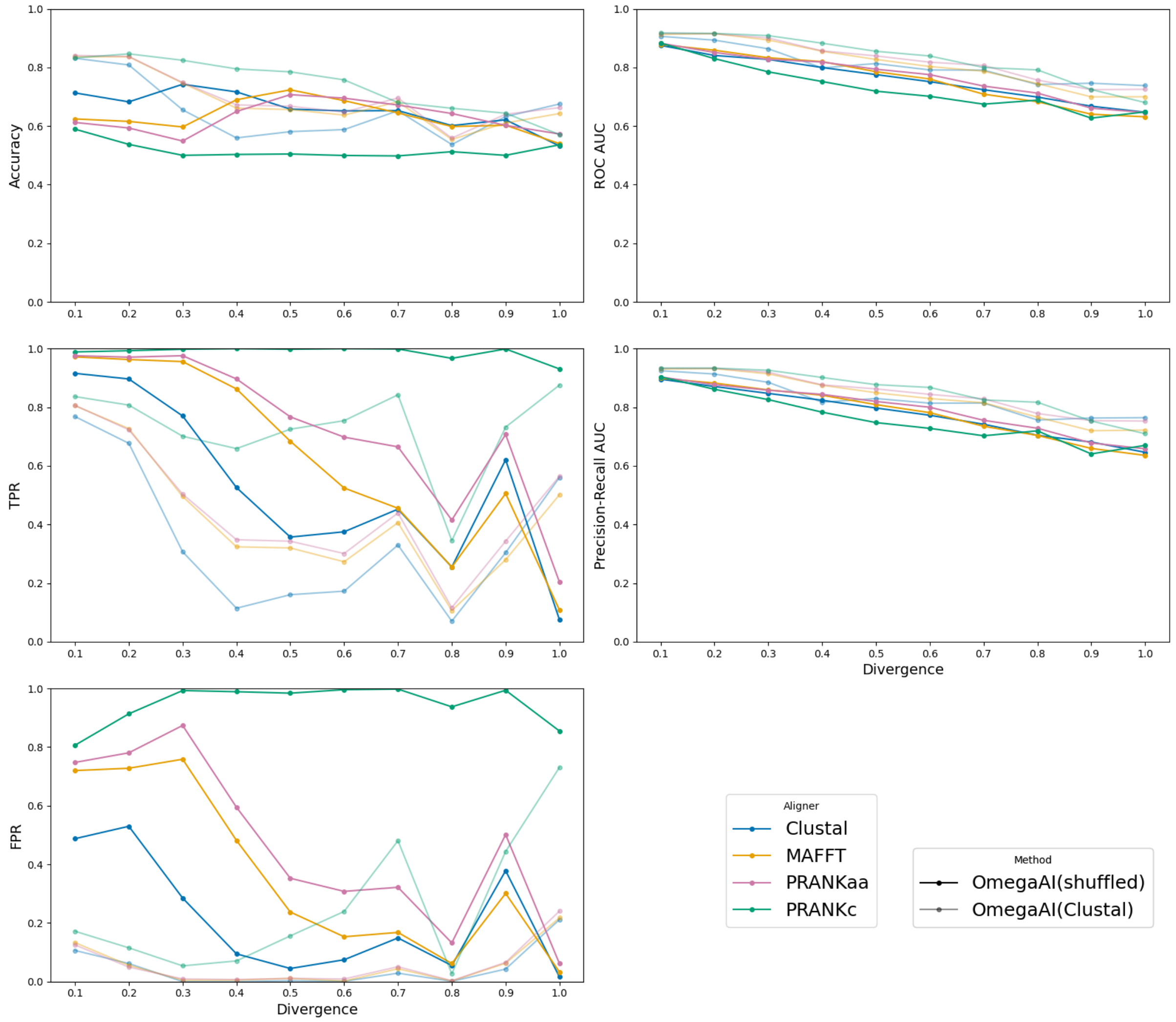


**Figure S 8. OmegaAI — shuffled vs. consistent MSA row ordering.** Various binary classifier performance metrics are presented to compare the two methods. Semi-transparent lines: results from the standard OmegaAI models that have been trained on Clustal alignments where row orderings are always consistent relative to the underlying simulation tree shown in Supplementary Fig. 1. Bold lines: results from OmegaAI models trained and tested on Clustal alignments where the row ordering has been randomised, meaning no information about the underlying tree is given to the models during training or testing. The divergence axis refers to the scaling of branches of the 8-taxa symmetric tree used for simulation. The baseline parameter for indel rate; 0.1, is used. These results show that the standard OmegaAI model leverages information about the tree in its learning and inference, and performance drops when it does not have access to this information.

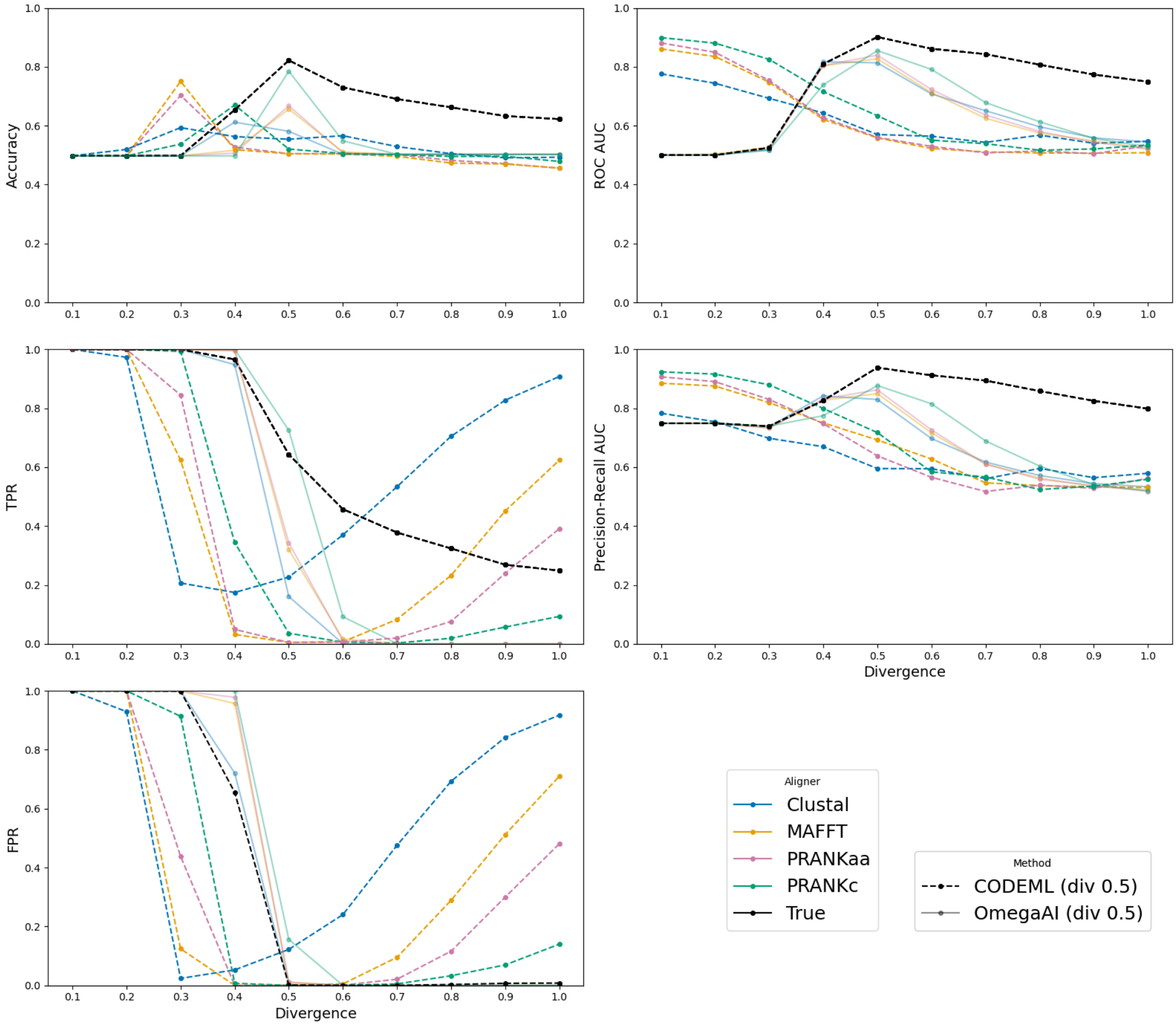


**Figure S 9. OmegaAI vs. CODEML — 0.5 divergence models.** Various binary classifier performance metrics are presented to compare the two methods. Semi-transparent lines show the OmegaAI model that has been trained on sequences simulated under the simulation tree with branches scaled to 0.5 substitutions per codon site per branch and aligned with Clustal. Dashed lines represent CODEML results, where CODEML has been forced to use the simulation tree with branches scaled to 0.5 during its free parameter value inference through maximum likelihood optimisation. Both OmegaAI and CODEML are tested with datasets across our divergence range, as indicated by the x-axis of each plot. Data at the divergence level of 0.5 is the only data for which the test data used has the same divergence level as is assumed by the methods. Colours represent the aligner used to align the test data. This analysis was undertaken to assess OmegaAI's ability to generalise to data that has evolved under different evolutionary rates compared to that which it was trained on. CODEML forced to use the 0.5 divergence tree is intended to be the analogous experiment in the maximum likelihood framework, for comparison. Both methods call positive selection for almost all data they are exposed to with divergence much less than 0.5. When tested using data with divergence greater than 0.5, OmegaAI tends to label almost all data with no positive selection. CODEML exhibits the opposite behaviour, where it overestimates positive selection and results in increasing TPR and FPR with increased divergence. The exception to this trend is CODEML applied to true alignments, where the trend looks more similar to that exhibited by OmegaAI, except CODEML is able to infer some true positives where OmegaAI struggles to do so.

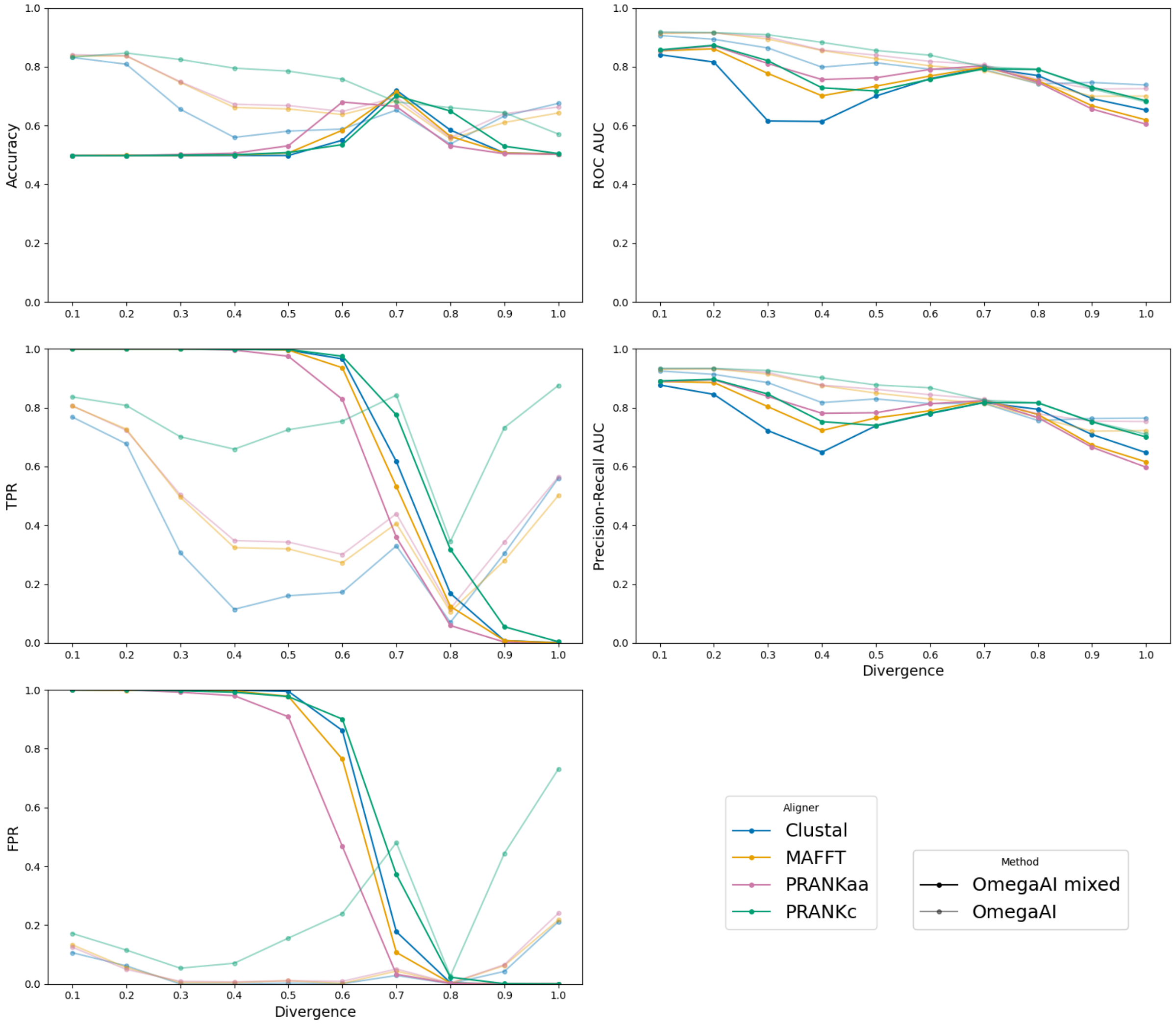


**Figure S 10. OmegaAI mixed model vs. individually trained OmegaAI models at each divergence.** Various binary classifier performance metrics are presented to compare the two methods. Semi-transparent lines show the results from individual OmegaAI models trained and tested for each divergence level in our divergence set, as seen in Figure 4. Models are trained using Clustal alignments. Solid lines show an OmegaAI model that has been trained on an equal number of trees from each divergence level in the set, with 1,000,000 in total. This model is then also tested across our divergence set. The analysis shows that a single OmegaAI model trained on different divergences is not immediately generalisable to all divergence testing scenarios because there is a thresholding issue, as indicated by low accuracy but high AUC values.

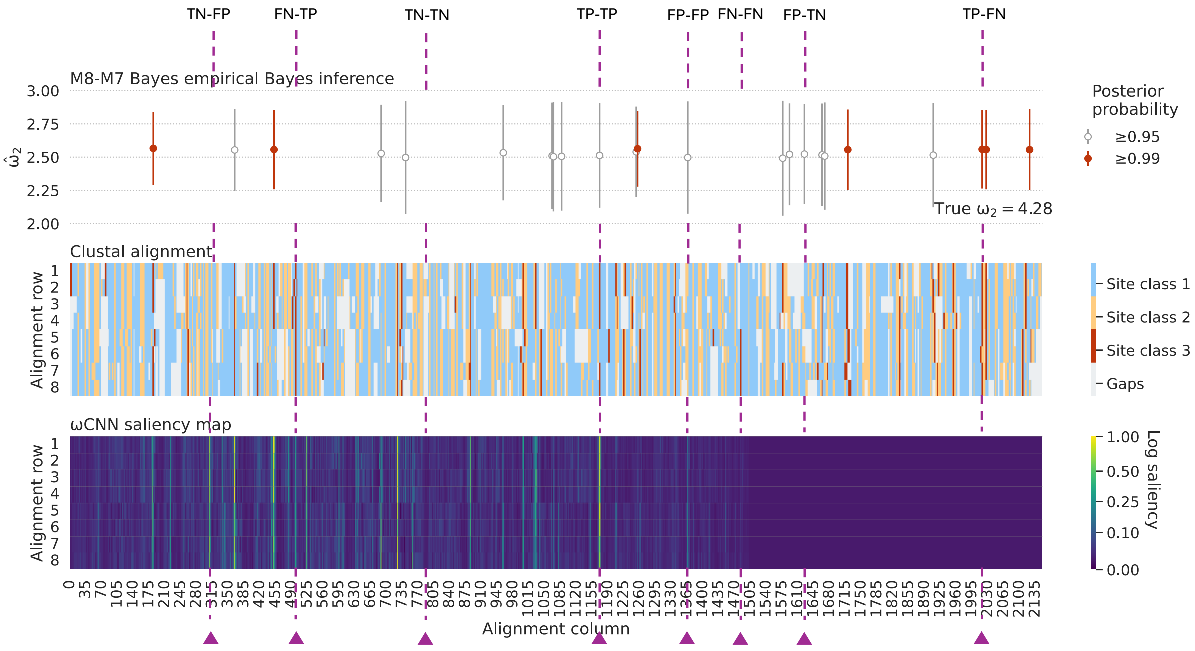


**Figure S 11. Comparing OmegaAI and CODEML inferences on sequences simulated under positive selection.** This plot and data is the same as seen in the main text Figure 9, but with further annotation. Vertical lines label examples of each of the 8 possible combinations of true/false positive/negative combinations when comparing OmegaAI and CODEML sitewise inferences of positive selection. However, we note that the OmegaAI negative inferences at the far right of the MSA are likely due to zero-padding during training (see Discussion). **Top panel:** CODEML sitewise predictions of positive selection. Following significant results from both M1a/M2a and M7/M8 LRTs (as described in Methods), an empirical Bayes approach is then used to calculate the posterior probability that each site is from a particular site class. Inferences of sites belonging to site classes with $\omega > 1$ with $p \geqslant 0.95$ or $p \geqslant 0.99$ are shown by grey and red bars, respectively. **Middle panel:** Colour coded Clustal alignment. Blue, yellow and red represent the three site classes described in Methods, with red indicating sites where $\omega > 1$. Grey indicates gaps in the alignment. **Bottom panel:** OmegaAI saliency map, computed as outlined in Methods.

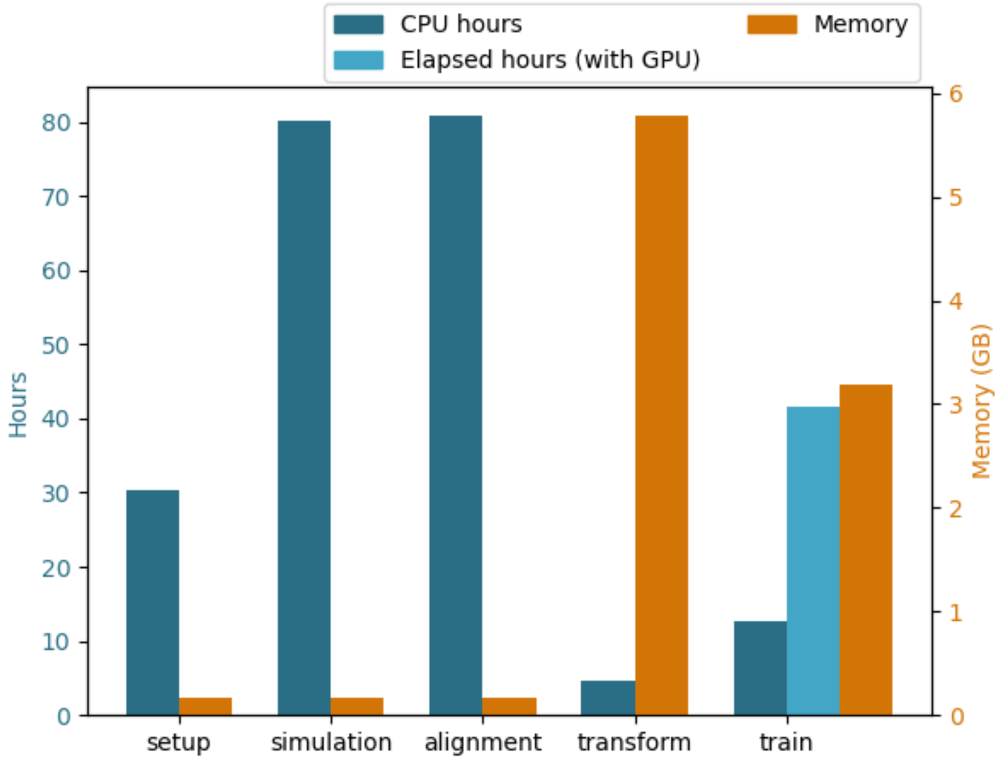


**Figure S 12. Benchmark resources for training OmegaAI.** CPU hours reflects an estimate of the time each step would take if run sequentially on a single CPU. In practice, most steps permit some amount of parallelisation when resources allow. Memory describes the maximum Resident Set Size (maxRSS) — the peak amount of physical memory actively used by a process during execution of each step. The workflow for training an OmegaAI CNN is split into the stages setup, simulation, alignment, data transformation (transform) and train. "Setup" describes processes such as directory creation, list creation for data chunking and various I/O. For the baseline model, 1,000,000 alignments are simulated using INDELible (Fletcher and Yang, 2009) and re-aligned using Clustal Omega (Sievers, Wilm, et al., 2011). Both processes are allocated 2 CPUs and 512MB of memory. "Transform" describes the process whereby MSAs are converted to TFRecord format. This step is given 1 CPU and 12GB of memory. Training involves a single CPU and a single GPU (NVIDIA Tesla V100 PCIe 32 GB). In this step, in addition to the total CPU time, we also include the elapsed hours to reflect the total time it takes to run the training step (c. 41.6 hours), for most of which the GPU is primarily utilised (GPU utilisation c. 75%).

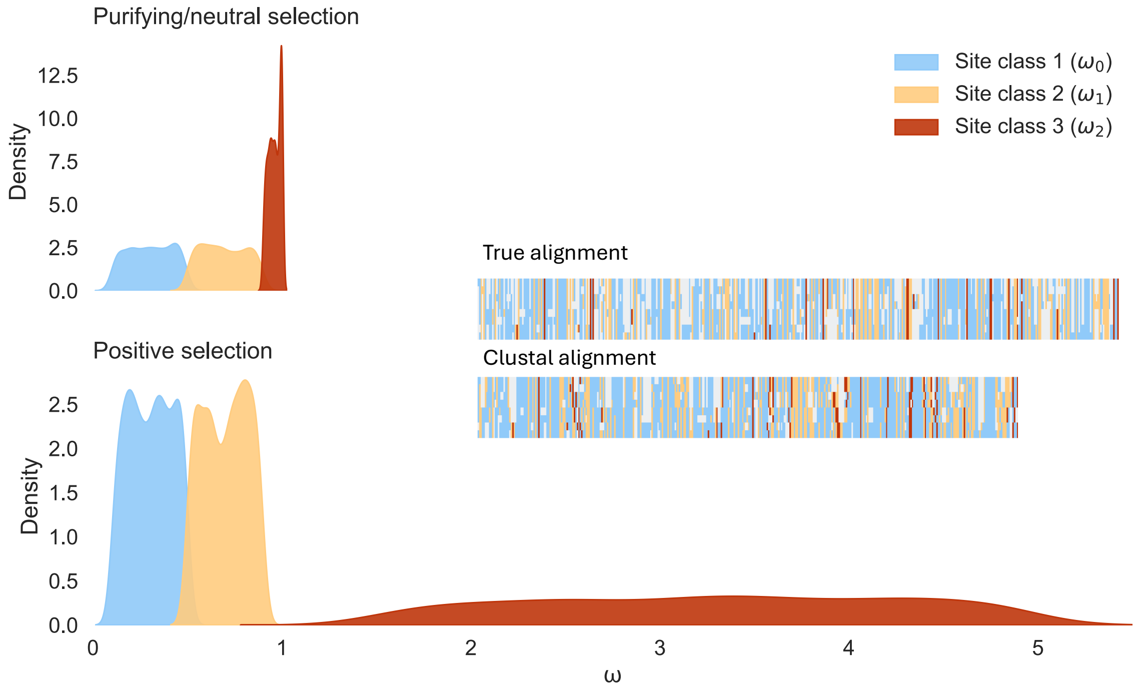


**Figure S 13. Distributions of $\omega$ values sampled for our "baseline" parameter simulations.** Kernel densities are shown to represent example distributions of $\omega$ values for the two simulation scenarios of purifying/neutral selection (top subplot) and positive selection (bottom subplot). These were created from empirically sampled $\omega$ values from the sampling distributions as defined in Methods. For purifying/neutral selection, $\omega$ values are always $\leqslant 1$. For positive selection, $\omega_2$ is always $\geqslant 1.5$ ($U(1.5, 5)$). An example is given of a colour-coded MSA from a true alignment and the same sequences re-aligned using Clustal. This illustrates how homologous codon sites become misaligned and the alignment becomes shorter due the mishandling of indels and over-alignment by Clustal.

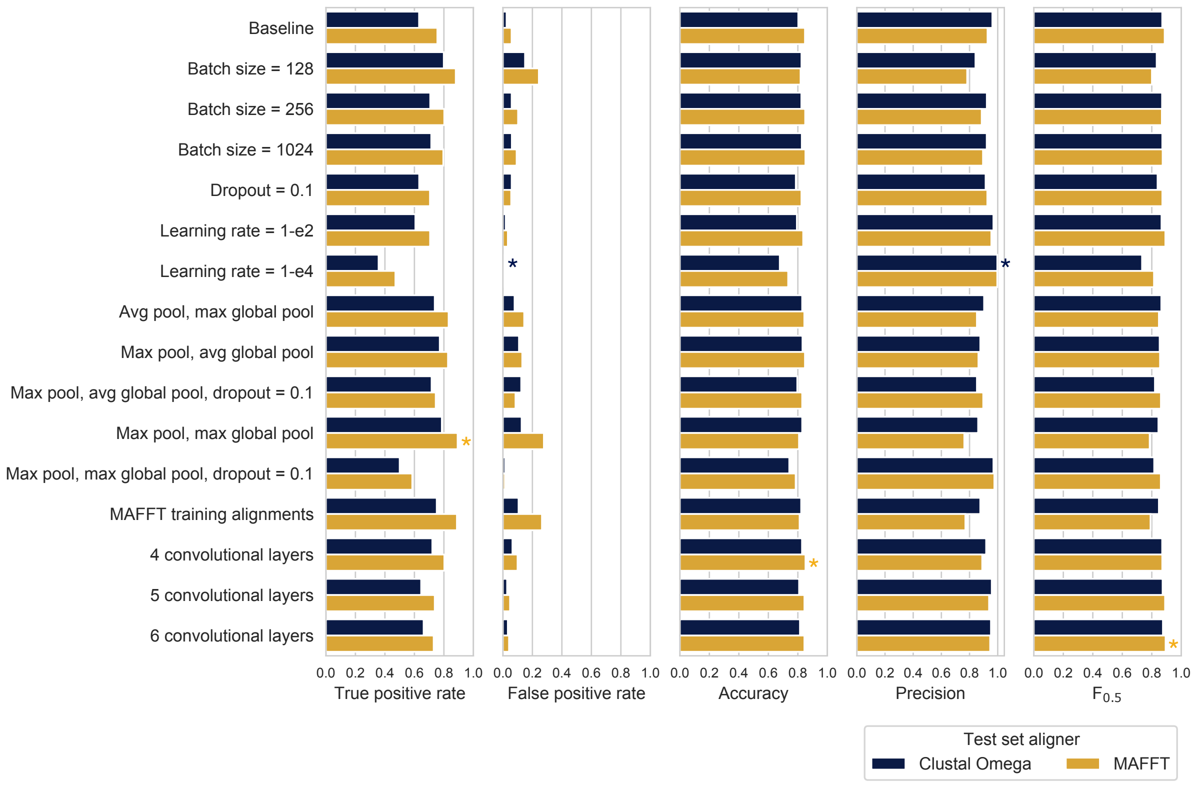


**Figure S 14. CNN performance with hyperparameter, training data, and architecture changes.** Starting with a "baseline" CNN architecture (top) and a "baseline" set of simulation parameters (see Methods), we trained the network for 50 epochs using 950,000 simulated alignments (re-aligned with Clustal Omega), and evaluated the performance of the trained network across a range of metrics (shown in separate subplots) by classifying a test set of 100,000 unseen Clustal Omega alignments, simulated using the same parameters as used for the "baseline" simulations. Then, for a selection of hyperparameters, including batch size, dropout probability, and learning rate, we re-trained the network and classified the same test set to compare performance. We also trained our network using MAFFT-aligned training data, different pooling operations, and with different numbers of convolutional layers. For each metric, an alignment colour-coded asterisk is shown beside the network/training data configuration which achieved the highest performance for that metric. As we were interested in limiting the number of false positives, we used the $F_{0.5}$ score as our primary metric for selecting the best performing network, for which the "6 convolutional layers" configuration achieved the best performance.

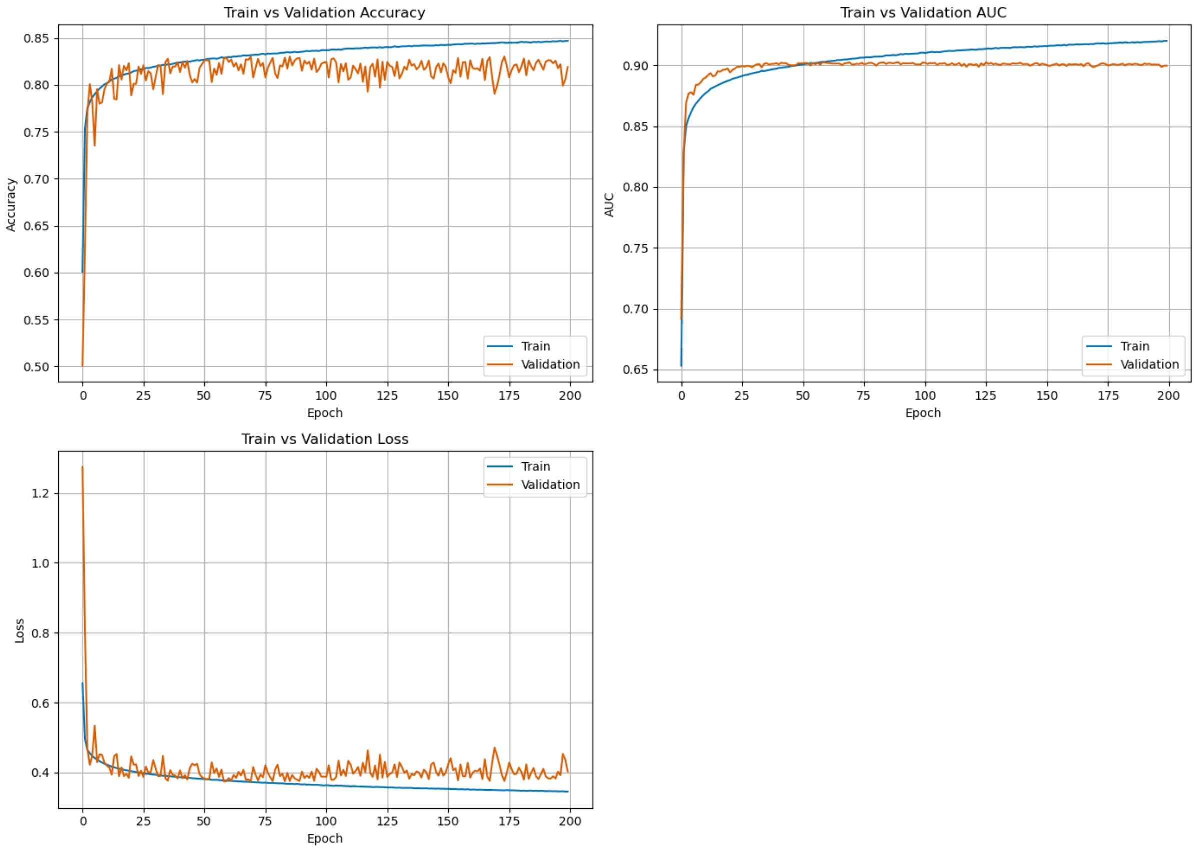


**Figure S 15. Training metrics over 200 epochs.** Accuracy, AUC and loss shown for both training (950,000 MSAs) and validation (50,000 MSAs) sets when training a baseline OmegaAI model over 200 epochs. Based on these analyses, the OmegaAI models in this study were trained for 50 epochs. This early stopping point prevents overfitting and avoids expending additional resources for negligible performance gains.

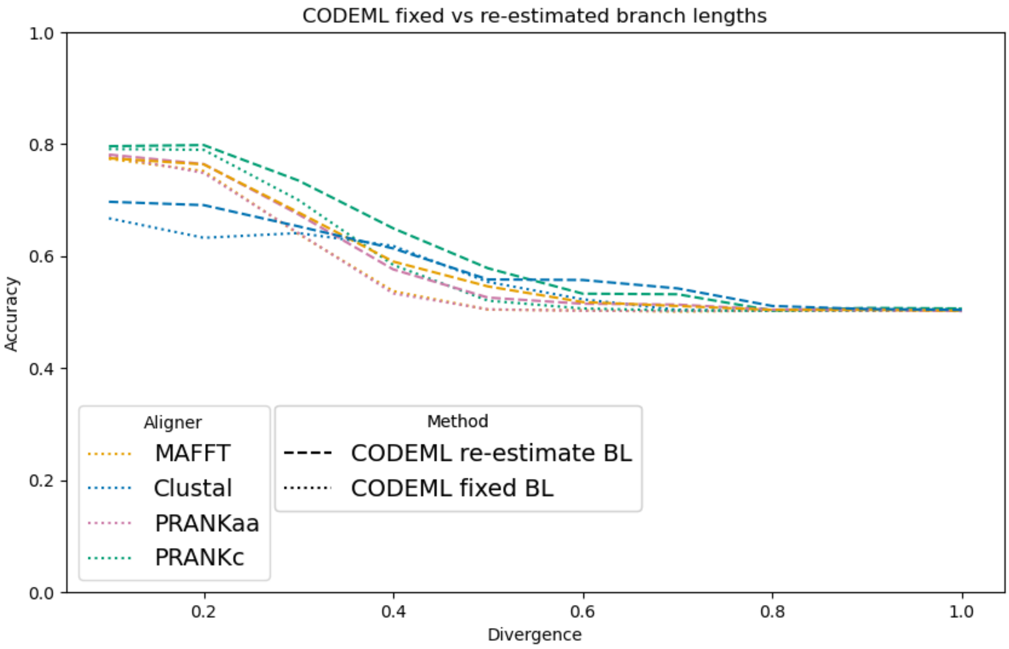


**Figure S 16. Comparing CODEML performance with fixed vs. re-estimated branch lengths.** CODEML was evaluated on sequences simulated under baseline conditions but with increasing divergence, for two scenarios, on four aligners. In both scenarios, CODEML is given the simulation topology of Supplementary Fig. 1 as the initial guide tree. In one scenario, during maximum likelihood estimation of parameters, CODEML is allowed to re-estimate branch lengths (results shown by dashed lines). In the other scenario the branch lengths are fixed (results shown by dotted lines). Across aligners and divergences, CODEML achieves higher accuracy when it is allowed to re-estimate branch lengths (as is common practice): this is the method presented throughout this work.
